# Evolution of glucuronoxylan side chain variability in vascular plants and the counter-adaptation of pathogenic cell-wall-degrading hydrolases

**DOI:** 10.1101/2024.01.23.575660

**Authors:** Li Yu, Louis F.L. Wilson, Oliver M. Terrett, Joel Wurman-Rodrich, Jan J. Lyczakowski, Xiaolan Yu, Kristian B.R.M. Krogh, Paul Dupree

**Author notes:** Corresponding author: Paul Dupree. These authors contributed equally to this paper (Li Yu, Louis F.L. Wilson, Oliver M. Terrett).

## Abstract

• Polysaccharide structural complexity not only influences cell wall strength and extensibility, but also hinders pathogenic and biotechnological attempts to saccharify the wall. In certain species and tissues, glucuronic acid side chains on xylan exhibit arabinopyranose or galactose decorations whose genetic and evolutionary basis is completely unknown, impeding efforts to understand their function and engineer wall digestibility.

• Genetics and polysaccharide profiling were used to identify the responsible loci in Arabidopsis and Eucalyptus from proposed candidates, while phylogenies uncovered a shared evolutionary origin. GH30-family *endo*-glucuronoxylanase activities were analysed by electrophoresis and their differing specificities were rationalised by phylogeny and structural analysis.

• The newly identified xylan arabinopyranosyltransferases comprise an overlooked subfamily in the GT47-A family of Golgi glycosyltransferases, previously assumed to comprise mainly xyloglucan galactosyltransferases, highlighting an unanticipated adaptation of both donor and acceptor specificities. Further neofunctionalisation has produced a Myrtaceae-specific xylan galactosyltransferase. Simultaneously, GH30 endo-glucuronoxylanases have convergently adapted to overcome these decorations, suggesting a role for these structures in defence. The differential expression of glucuronoxylan-modifying genes across Eucalyptus tissues, however, hints at further functions.

• Our results demonstrate the rapid adaptability of biosynthetic and degradative carbohydrate-active enzyme activities, providing insight into a plant-pathogen arms race and facilitating plant cell wall biotechnological utilisation.

## Introduction

Surrounding every plant cell, the cell wall is an extracellular matrix that generates turgor pressure and affords protection from abiotic and biotic stresses. Cell walls are composed of a complex combination of polysaccharides, phenolic polymers, and proteins; however, despite a universal and fundamental role in cell biology, their structural compositions have diverged dramatically and rapidly throughout evolution (Burton *et al*., 2010). Given the cell wall’s importance both for morphology and as a complex physical and metabolic barrier to pathogens, this diversity likely results not only from adaptation to different physical environments, but also from a complex ongoing arms race between plants and their enemies (Malinovsky *et al*., 2014).

Forming a common hemicellulosic component of plant cell walls, xylan polysaccharides are thought to bind to and thus chaperone cellulose microfibrils (Busse-Wicher *et al*., 2014; Simmons *et al*., 2016) as well as mediating cross-links to other cell wall components (Terrett & Dupree, 2019; Tryfona *et al*., 2023). Xylans vary in structure between different species and tissues, but invariably they possess a backbone of β1,4-linked xylosyl residues (Scheller & Ulvskov, 2010). In Arabidopsis and other eudicots, the xylan backbone is typically decorated with 2/3-linked acetate and α1,2-linked glucuronic acid (GlcA), which is often 4-*O*-methylated. The spacing of these decorations along the xylan backbone is critical for regulating its interaction with cellulose microfibrils (Simmons *et al*., 2016; Grantham *et al*., 2017; Tryfona *et al*., 2023). While most xylan in Arabidopsis is present in the secondary cell wall (SCW), it also makes up part of the primary cell wall (PCW) (Mortimer *et al*., 2015). Although the function of PCW xylan has not been determined, plants appear to allocate dedicated machinery to synthesise it: whereas the SCW xylan backbone is synthesised by IRX9, IRX10, and IRX14/14L, PCW xylan is synthesised specifically by IRX9L, IRX10L, and IRX14 (Mortimer *et al*., 2015; Anders *et al*., 2022). Furthermore, the pattern of GlcA decoration differs substantially between PCW and SCW xylan. SCW xylan glucuronidation is achieved by two enzymes: GUX1, which produces an even-spaced pattern, and GUX2, which produces a more random pattern of GlcA side chains (Bromley *et al*., 2013). In contrast, PCW xylan is glucuronidated consistently at every sixth backbone xylosyl residue by GUX3 (Mortimer *et al*., 2015). Notably, in at least some PCW-rich Arabidopsis tissues, the GlcA moieties of PCW xylan can be further decorated with a pentose that has been assigned as a putative α1,2-linked L-arabinopyranose (Ara*p*) (Chong *et al*., 2015; Mortimer *et al*., 2015). The glycosyltransferase responsible for this decoration has not yet been identified, hindering investigations into the functions of these Ara*p*-α1,2-GlcA-α1,2-(U^Ara*p*^) disaccharide sidechains.

Prior to its discovery in Arabidopsis, decoration of xylan GlcA residues with additional sugars was previously observed for several species in the *Eucalyptus* genus of the Myrtaceae family (Shatalov *et al*., 1999; Evtuguin *et al*., 2003; Pinto *et al*., 2005). Interestingly, further experiments clearly showed that the GlcA residues in Eucalyptus xylan can not only be substituted with α1,2-linked L-Ara*p* (as thought to be the case in Arabidopsis) but also with a sterically similar moiety: β1,2-linked D-galactose (Gal) (Togashi *et al*., 2009; Pena *et al*., 2016). Furthermore, the U^Ara*p*^ sidechain has also been shown to be widely present in xylans from non-commelinid monocots (Pena *et al*., 2016). However, whether the Gal-β1,2-GlcA-α1,2-(U^Gal^) sidechain is present in plants outside the *Eucalyptus* genus is unknown.

Disaccharide side chains terminating in 2-linked sugars, particularly β1,2-linked galactose, can also be found decorating other PCW hemicelluloses, namely xyloglucan and β-galactoglucomannan (β-GGM). Arabidopsis mutants lacking the galactosyl residues of xyloglucan’s Gal-β1,2-Xyl-α1,6-side chains exhibit stunted growth (Tamura *et al*., 2005), while further removal of the equivalent residue in β-GGM’s Gal-β1,2-Gal-α1,6-side chains exacerbates the phenotype (Yu *et al*., 2022). These side chain structures may influence the ability of these hemicelluloses to achieve their presumed primary function in chaperoning and cross-linking cellulose microfibrils. However, as well as contributing to the material properties of the cell wall, PCW hemicelluloses pose an obstacle to invading pathogens. Pathogens must therefore secrete cell wall-degrading glycosyl hydrolases (GHs) in order to gain access to the cell, and the added structural complexity afforded by cell wall polysaccharide decorations likely increase the difficulty in doing so.

In particular, to overcome the barrier posed by xylan, fungal and bacterial pathogens typically secrete a variety of xylanolytic enzymes. Xylanase activity is thought to be an important part of infection by disease-causing microbes (Bellincampi *et al*., 2014), and cereals such as wheat produce xylanase-inhibitory proteins in response to fungal infection (Chmelová *et al*., 2019). Xylan degradation begins with backbone cleavage by *endo*-xylanases, the best studied being from CAZy families GH5, 8, 10, 11, and 30 (Paes *et al*., 2012). Enzymes from GH8, 10, and 11 tend to cut preferentially at unsubstituted regions of the xylan backbone. Such enzymes are impeded by side chain decorations, which also require later deconstruction by dedicated *exo*-glycosidases (terminal GlcA residues are removed by α-glucuronidases from families GH67 and GH115, for instance). However, some *endo*-xylanases have specifically evolved to recognise certain side chains: most characterised GH30 xylanases require a GlcA-substituted xylosyl residue at the −2 subsite, while GH5 xylanases can be arabinoxylan-specific. Nevertheless, binding of such enzymes is inhibited by incompatible or clustered side chains. For these reasons, additional structural complexity provides increased resistance to degradation, and plants are presumably under a constant selection pressure to innovate new and more complex xylan structures. An example of such adaptation may lie in corn bran xylan, whose unique complexity provides exceptional recalcitrance to enzymatic digestion (Rogowski *et al*., 2015). In turn, pathogens are under a selection pressure to develop hydrolases that can accommodate these more complex xylan structures (Beri *et al*., 2020).

Investigating the true functions of such cell wall structural features, as well as the evolutionary forces behind their continuing diversification, is impossible without knowing the relevant biosynthetic genes. In the case of xylan, the genes necessary for arabinopyranosylating and galactosylating glucuronic acid side chains are still unknown. Here we report that a family of mainly xyloglucan-modifying glycosyltransferases have been remodelled through evolution for this purpose, requiring reprogramming of both donor sugar and backbone acceptor specificities to generate the new xylan-arabinopyranosylating activity. We characterise the activities of three family members through heterologous expression in Arabidopsis, and show that, moreover, this family has itself undergone even more recent neofunctionalisation in the Myrtaceae family to produce xylan galactosyltransferases, explaining the previously observed xylan structures in Eucalyptus. Furthermore, we show that the relative expression of xylan arabinopyranosyltransferases and galactosyltransferases across Eucalyptus tissues defines two contrasting distributions of U^Ara*p*^ and U^Gal^ structures, suggesting a potential functional role in specialised cell wall architectures. Finally, we show not only that the decoration of GlcA with Ara*p*/Gal impedes the action of some GH30 glucuronoxylanases (suggesting a role in passive defence) but also that a minority of GH30 family members have convergently acquired amino acid substitutions to accommodate such structures. As well as providing insight into the synthesis and function of PCW xylan and the arms race between plants and pathogens, our findings are therefore of benefit in improving industrial hemicellulosic biomass degradation.

## Materials and Methods

### Plant Material

All Arabidopsis (*Arabidopsis thaliana* _(L.) Heynh._) T-DNA insertion mutant lines were in the Col-0 ecotype. *At1g68470* (*Atgt17/xapt1*): WiscDsLox437H04 (*xapt1-1*), *at2g29040* (*Atgt11*): Salk_108349, *at2g32750* (*Atgt12*): Salk_057536, *at2g32740* (*Atgt13*): Salk_053593C, *at4g22580* (*Atgt19*): Salk_075020 and *at5g41250*: Salk_085915 (*Atgt20*) were obtained from the Nottingham Arabidopsis Stock Centre (NASC, Nottingham, UK). Homozygous lines were isolated by PCR. The *irx9l* (SALK_037323) and *mur3-3* mutants have been described previously (Wu *et al*., 2010; *Kong et al.*, 2015). All the primers used are shown on Table S1. Arabidopsis young stem was harvested at 30 days and bottom stem was harvested at 8 weeks.

Actively growing *Eucalyptus dalrympleana* _Maiden_ branch and leaves were harvested at Jesus College, University of Cambridge in April 2018. Young, actively growing *Myrtus communis* _L._ and *Metrosideros excelsa* _Sol. ex Gaertn_. branchlets were harvested at Cambridge University Botanic Garden, and *Plinia cauliflora* _(Mart.) Kausel_, *Psidium guajava* _L._, and *Myrcianthes pungens* _(O.Berg) D.Legrand_ material was a gift of Pedro Araújo (University of Campinas). Combined phloem and periderm cortex material was separated from xylem/pith by peeling or scraping. After collection, all the materials were put into 96% ethanol. For all samples, alcohol insoluble residue (AIR) was prepared as described previously (Goubet *et al*., 2009).

### Plant Growth and Arabidopsis transformation

Arabidopsis plants were grown on a 9:1 mix of Levington M3 compost and vermiculite and grown at 21°C under a 16 h/8 h photoperiod. Constructs used for *A. thaliana* Xylan ArabinoPyranosylTransferase 1 (*At*XAPT1), *Eucalyptus grandis* _W.Hill ex Maiden_ (*Eg*XAPT), and Xylan GalactoPyranosylTransferase (*Eg*XLPT) overexpression in *xapt1* and WT plants were prepared using OpenPlant common syntax GoldenGate assembly (Patron *et al*., 2015). *AtXAPT1* (*At1g68470*), *EgXAPT* (*Eucgr.D00738*; GenBank: XM_010054097.3), and *EgXLPT* (*Eucgr.H00343*; GenBank: XM_010070979.3) coding sequences and the *CESA3* and *IRX3* promoter sequences were synthesised *de novo* by IDT or GeneWiz after removal of *BsaI* and *BpiI* restriction sites. Arabidopsis transformation was performed using the floral dip method as previously described (Clough & Bent, 1998).

### Phylogenetic and bioinformatics analysis

For the XAPT phylogeny, XAPT orthologues (GT47-A_V_) from a wide range of tracheophyte species were extracted from our recent phylogeny of GT47-A enzymes (Yu *et al*., 2022). To obtain further XAPT orthologues from plants in the Myrtales order, the genomes of *Metrosideros polymorpha* (Izuno *et al*., 2016; Izuno *et al*., 2019), *Rhodamnia argantea*, *Syzygium oleosum*, and *Punica granatum* (https://www.ncbi.nlm.nih.gov/genome/) and the transcriptome of *Oenothera rosea* (Carpenter et al., 2019; One Thousand Plant Transcriptomes, 2019), were searched using TBLASTN (Altschul *et al*., 1990) using *At*XAPT1 as a query with a cut-off at *E* = 1 × 10^-125^. Sequences were aligned using MUSCLE (Edgar, 2004a; Edgar, 2004b) and truncated to the GT47 domain. ProtTest3 (Darriba *et al*., 2011) was then used to select an appropriate substitution model, and a phylogeny was calculated using RAxML (Stamatakis, 2014) with 100 rapid bootstraps. For the main text figure, the tree was then trimmed to species of interest using ape 5.0 (Paradis & Schliep, 2019).

For the GH30 subfamily 8 tree, CAZy sequences were downloaded from the dbCAN2 server (http://bcb.unl.edu/dbCAN2/) (Zhang *et al*., 2018). Using bespoke Python scripts invoking HMMER (Eddy, 2011), GH30_8 domain-containing sequences were extracted and trimmed to the GH30 domain. Sequences with a GH30 domain sequence shorter than 200 residues were omitted. Extremely similar sequences were removed using CD-HIT (Li & Godzik, 2006; Fu *et al*., 2012) with a 99 % similarity threshold. The remaining sequences were aligned using MAFFT (Katoh & Standley, 2013); ProtTest3 was then used to select an appropriate substitution model, and a phylogeny was calculated using RAxML with 100 rapid bootstraps.

### Genomic DNA extraction and sequencing of native *XAPT* and *XLPT* loci

*E. dalrympleana* leaves were ground in liquid nitrogen and genomic DNA was extracted as previously described (James *et al*., 2001), supplementing the extraction buffer with 3 % polyvinylpyrrolidone. PCR amplification was achieved using Q5 High-Fidelity DNA Polymerase (Biolab); primers are listed in Table S1. PCR products were purified by gel extraction and isopropanol precipitation before Sanger sequencing.

### Enzyme Digestions

Soluble hemicelluloses were prepared from AIR as previously described (Lyczakowski *et al*., 2017). For Arabidopsis bottom stem and *E. dalrympleana* xylan analysis, 0.5 mg AIR was used for one reaction. For Arabidopsis young stem, 1 mg AIR was used for one reaction.

Soluble hemicellulose fraction was digested with an excess of Xyn11A from *Neocallimastix patriciarum* (*Np*Xyn11A, Genbank: CAA46498.1; from Megazyme) in 50 mM ammonium acetate, pH 6.0 at 30°C overnight.

Alternatively, xylan was digested with GH30 xylanases with a 70 μg ml^-1^ concentration from different bacteria at 25°C for 30 min in 50 mM ammonium acetate, pH 6.0. The GH30 xylanases included: XynC from *Bacteroides ovatus (Bo*XynC, GenBank: ALJ48333.1; a gift of Harry Gilbert, Newcastle University); Xyn30A from *Hungateiclostridium thermocellum* (*Ct*Xyn30A, GenBank: ABN54208.1; from NZYTech). XynA from *Dickeya chrysanthemi* strain D1 (*Ec*_D_Xyn30A, GenBank: AAB53151.1), XynA from *Dickeya chrysanthemi* strain P860219 (*Ec*_P_Xyn30A, GenBank: AAL16415.1), and the two mutagenized enzymes *Ec*_D_Xyn30A_Y255L_ and *Ec*_P_Xyn30A_L255Y_ were provided by Novozymes A/S. For mass spectrometry and galactosidase sensitivity experiments, xylanase products were then digested overnight with 10 μL of 1 mg ml^-1^ *B. ovatus* GH115 glucuronidase (*Bo*Agu115A, GenBank: EDO10816.1; a gift of Harry Gilbert, Newcastle University) to remove the non-decorated GlcA side chains.

For the galactosidase sensitivity assay, the xylanase products were passed through a 3-kDa Nanosep centrifugal device. Next, ethanol was added to a final concentration to 65% before incubation at −20°C for 2 h. The samples were then centrifuged at 12,000g for 10 min; the supernatant was subsequently dried 60°C *in vacuo* and resuspended in 200 μl of 50 mM ammonium acetate, pH 6.0. To this, 0.5 μl of 4 kU ml^-1^ GH35 β-galactosidase from *Aspergillus niger* (Megazyme) was added and incubated at 37°C for 18 h. Samples were then dried at 60°C *in vacuo*.

### Oligosaccharides Fingerprint Analysis by PACE

Samples and (Xyl)_1–6_ standards (Megazyme, Ireland) were derivatized with 8-aminonapthalene-1,3,6-tresulphonic acid (ANTS; Invitrogen) as described previously (Goubet et al., 2002). After drying, the samples were resuspended in 100 μl of 3 M urea, of which 2 μl was loaded onto the PACE gels. The samples were run and visualized as described previously (Goubet *et al*., 2009).

### Preparation of Oligosaccharides for MS

Following the enzyme digestion, released peptides and enzymes were removed using reverse-phase Sep-Pak C18 cartridges (Waters) as previously described. Oligosaccharides were reductively aminated with 2-aminobenzoic acid (2-AA) and cleaned using a GlycoClean S cartridge (ProZyme) as previously described (Tryfona & Stephens, 2010).

## Results

### At1g68470 encodes XYLAN ARABINOPYRANOSYLTRANSFERASE 1 (XAPT1) in Arabidopsis

Despite an almost-complete genetic understanding of xylan biosynthesis, the gene(s) responsible for arabinopyranosylating and galactosylating glucuronic acid side chains remain unknown, thwarting efforts to determine their function and manipulate xylan structure. To remedy this, we sought to obtain candidate arabinopyranosyltransferase mutants in Arabidopsis, which normally exhibits putative U^Ara*p*^ (Ara*p*-α1,2-GlcA-α1,2-) xylan side chains in its primary cell wall. However, according to the CAZy database (Drula *et al*., 2022), the Arabidopsis genome encodes 558 glycosyltransferases (GTs) distributed amongst at least 43 GT families, of which only a minority are functionally characterised; too large a number to be screened experimentally. Instead, we attempted to predict likely candidates through biochemical plausibility. Firstly, we observed that the α1,2-linked arabinopyranosyl moiety of the U^Ara*p*^ structure is sterically almost identical to the β1,2-linked galactosyl moiety of the Eucalyptus U^Gal^ structure (Gal-β1,2-GlcA-α1,2-), differing only in the presence or absence of a C6 hydroxymethyl group. Therefore, we hypothesised that the two corresponding glycosyltransferases may be closely related. Secondly, owing to our recent work on xyloglucan and β-GGM synthesis, we observed that all subfamily A members of the CAZy GT47 family (GT47-A) characterised so far exhibit an inverting mechanism and affix the donor sugar, most commonly galactose, using a 1,2-glycosidic linkage (MUR3, for instance, creates the Gal-β1,2-Xyl-α1,6-‘L’ side chain of xyloglucan). Furthermore, all known acceptors are α-linked monosaccharide decorations of a β1,4-linked hemicellulose backbone. These intriguing similarities to the theoretical activities of xylan arabinopyranosyltransferases and galactosyltransferases led us to hypothesise that the responsible genes may reside in the same family.

We recently calculated a large-scale phylogeny of the GT47-A subfamily in plants (Yu *et al*., 2022), which revealed that the subfamily consists of at least seven subclades (GT47-A_I–VII_). Importantly, xyloglucan-modifying activities are spread almost universally throughout these clades, which has always given the impression that all unidentified members share this activity. Nevertheless, several Arabidopsis members—as well as two subclades (A_IV_ and A_V_)—are completely uncharacterised; hence, we hypothesised that these previously overlooked enzymes could harbour unanticipated xylan-modifying activities. To test this hypothesis, we attempted to detect the loss of the PCW xylan U^Ara*p*^ structure in T-DNA knockout lines for the six remaining Arabidopsis GT47-A subclade members with unknown functions (*At5g41250/AtGT20* from A_II_, *At4g22580/AtGT19* from A_IV_, *At1g68470/AtGT17* from A_V_, and *At2g29040/AtGT11*, *At2g32750/AtGT12*, and *At2g32740/AtGT13* from A_VII_). *At2g29040*/*AtGT11* has since been implicated in galactosylation of pollen tube xyloglucan (Wei *et al*., 2021). We used the *irx9l* mutant, which lacks PCW-type xylan (and therefore does not exhibit any U^Ara*p*^ structure), as a control (Mortimer *et al*., 2015). AIR from the young stems of these mutants was incubated with alkali to solubilize the hemicelluloses, and then digested with *Np*Xyn11A *endo*-xylanase. The resultant oligosaccharides were analysed by PACE. As previously reported (Mortimer *et al*., 2015), wild-type (WT) young stem yielded a series of products including xylose (X), xylobiose (X_2_), aldopentaouronic acid with (XU^m^XX) or without (XUXX) 4-O-methyl group on the GlcA (generically termed ‘XU^[m]^XX’), and xylopentaose substituted with the putative Ara*p*-GlcA-disaccharide (XU^Ara*p*^XXX). In contrast, digestion of material from the *irx9l* mutant released xylose, xylobiose (X_2_), and XU^[m]^XX, but not XU^Ara*p*^XXX. Interestingly, although we could detect XU^Ara*p*^XXX in five of the six GT47-A mutants, this product was also conspicuously absent for *at1g68470* (A_V_), suggesting that xylan in this mutant lacks the U^Ara*p*^ decoration (Fig. 1a, S1). To help confirm this, WT young stem was hydrolysed with the GH30-family xylanase Xyn30A from *Bacteroides ovatus* (*Bo*XynC), which requires a GlcA substitution at its −2 subsite. As previously described (Mortimer *et al*., 2015), digestion of WT material with *Bo*XynC produced XXXXU^[m]^X (U_1_^[m]^X_6_) and XXXXU^Arap^X (U_1_^Ara*p*^X_6_) as major products, in accordance with the consistent GlcA spacing of PCW xylan. In contrast, digestion of material from the *at1g68470* mutant released U_1_ ^[m]^X_6_ but not the U ^Ara*p*^X oligosaccharide (Fig. 1b). These results suggest that, unexpectedly, *At1g68470* is required for the synthesis of the putative Ara*p*-GlcA-disaccharide substitution on Arabidopsis PCW xylan.

**Figure 1.**
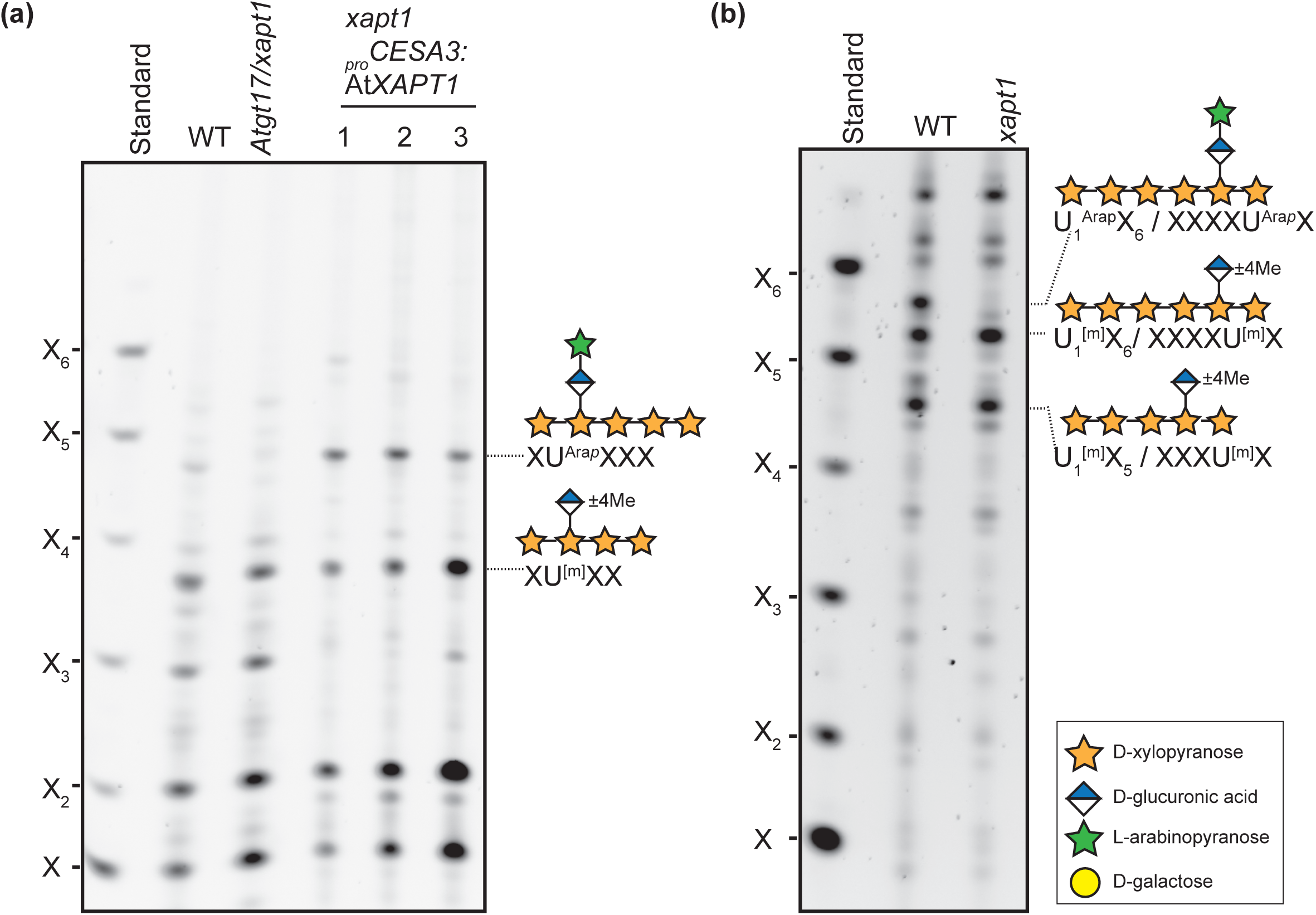
AT1G68470/XAPT1 from GT47 subclade A is a functional xylan arabinopyranosyltransferase. (a), PACE analysis of *Np*Xyn11A xylanase digests of alkali-extracted xylan from AIR of WT, the *xapt1* T-DNA insertion mutant, and three independent transgenic lines expressing *XAPT1* under the strong primary wall-specific promoter of *CESA3* in the *xapt1* background. (b), PACE analysis of *Bo*XynC xylanase digests of WT and *xapt1* mutant young stem. Xylo-oligosaccharide standard: X–X_6_.

To further support our conclusion, we complemented the *at1g68470* mutant with *At1g68470* under the control of the PCW-synthesis-specific *CESA3* promoter. AIR from young stems was digested by xylanase *Np*Xyn11A and analysed by PACE. In all three independent transgenic lines, the XU^Ara*p*^XXX product was detected at substantially higher levels than in WT (Fig. 1a). Based on these data, and the previous arabinopyranose assignment, we concluded that *At1g68470* likely encodes a xylan-specific arabinopyranosyltransferase that catalyses the formation of an Ara*p*-α1,2-GlcA linkage. We therefore propose that *At1g68470* is named ***X****YLAN **A**RABINO**P**YRANOSYL**T**RANSFERASE 1* (*XAPT1*).

### XAPT underwent gene duplication in the Myrtales

Having successfully predicted the identity of *XAPT1* in Arabidopsis, we also investigated our hypothesis that xylan galactosyltransferases could be closely related. Importantly, whereas the U^Ara*p*^ xylan branch structure has been reported in a wide range of species, including Arabidopsis, *Eucalyptus* spp., and non-commelinid monocots, the U^Gal^ structure has so far only been detected in *Eucalyptus* spp. (Togashi *et al*., 2009; Pena *et al*., 2016). This suggests that xylan galactosyltransferases have in fact evolved more recently than XAPTs. We hypothesised that: *a)* XAPT1’s parent subclade, GT47-A_V_, comprises (mainly) xylan arabinopyranosyltransferases, and *b)* that xylan galactosyltransferases also derive from this subclade. To explore this idea, we constructed a phylogeny of XAPT homologues from 56 species of vascular plant, including *Eucalyptus grandis*, three fellow members of the Myrtaceae family (*Metrosideros polymorpha*, *Syzygium oleosum*, and *Rhodamnia argantea*), and two members of the wider Myrtales order (*Punica granatum* and *Oenothera rosea*). Fig. 2 presents the rooted phylogeny, which has been trimmed for brevity (the full phylogeny can be found in Fig. S2). We found that most of the plant genomes—including *P. granatum* and *O. rosea*—exhibited a single gene (or cluster of near-identical copies) in this subclade. In contrast, all four Myrtaceae genomes exhibited two paralogues that appear to have arisen from a single, recent duplication event that likely occurred in the Myrtales order. In the case of *E. grandis*, these two paralogues correspond to the genes *Eucgr.D00738* and *Eucgr.H00343*. Hence, we hypothesised that Eucgr.D00738 (*Eg*XAPT, hereafter) and Eucgr.H00343 (*Eg*XLPT) could each constitute either a xylan GlcA arabinopyranosyltransferase or potentially a xylan GlcA galactosyltransferase.

**Figure 2.**
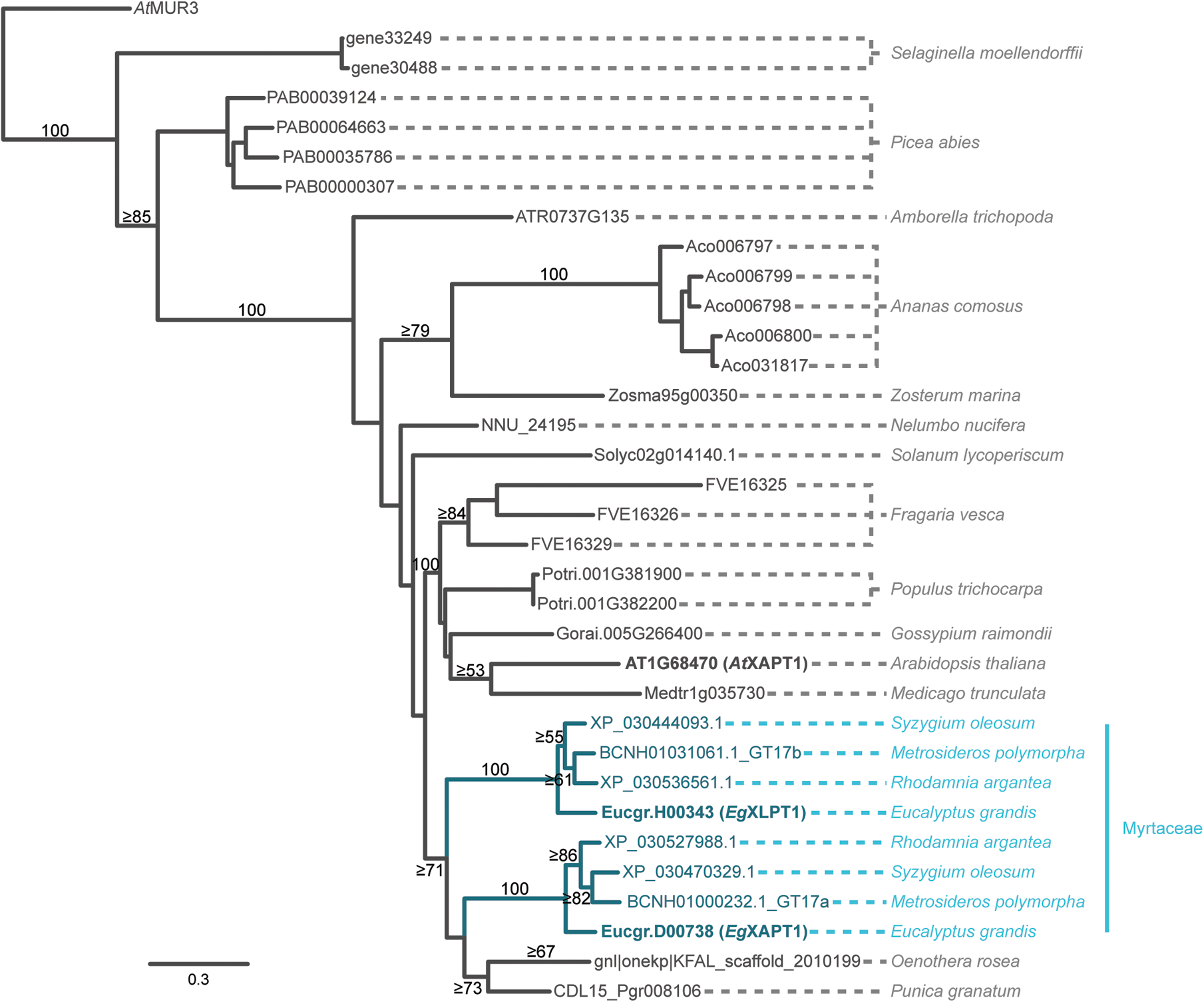
Trimmed phylogenetic tree of the XAPT subclade of GT47-A_V_. Sequences grouped in the XAPT subclade of our previous GT47-A tree (Yu et al., 2022) were supplemented with HMMER and TBLASTN hits for XAPT-like sequences in Myrtales genomes. Following alignment, the phylogeny was inferred using RAxML. For rooting, *At*MUR3 was included as the outgroup. Due to the large number of sequences (106), we show only an abridged form of the tree here (displaying the more pertinent sequences), hence bootstrap values (from 100 pseudoreplicates) may underestimate the confidence of the splits shown. The full phylogeny can be viewed in Fig. S3.

### Eucgr.D00738 and Eucgr.H00343 encode EgXAPT and EgXLPT

To investigate the activities of the two Myrtaceae XAPT-related paralogues, we expressed *Eg*XAPT and *Eg*XLPT under the strong, PCW-synthesis-specific *CESA3* promoter in the *xapt1* Arabidopsis mutant. Xylan digestion with *Np*Xyn11A xylanase revealed that expression of either gene could restore the presence of disaccharide side chain structures (Fig. 3a), suggesting that both proteins are active and catalyse the formation of Ara*p*-α1,2-GlcA or a similar linkage on xylan.

**Figure 3.**
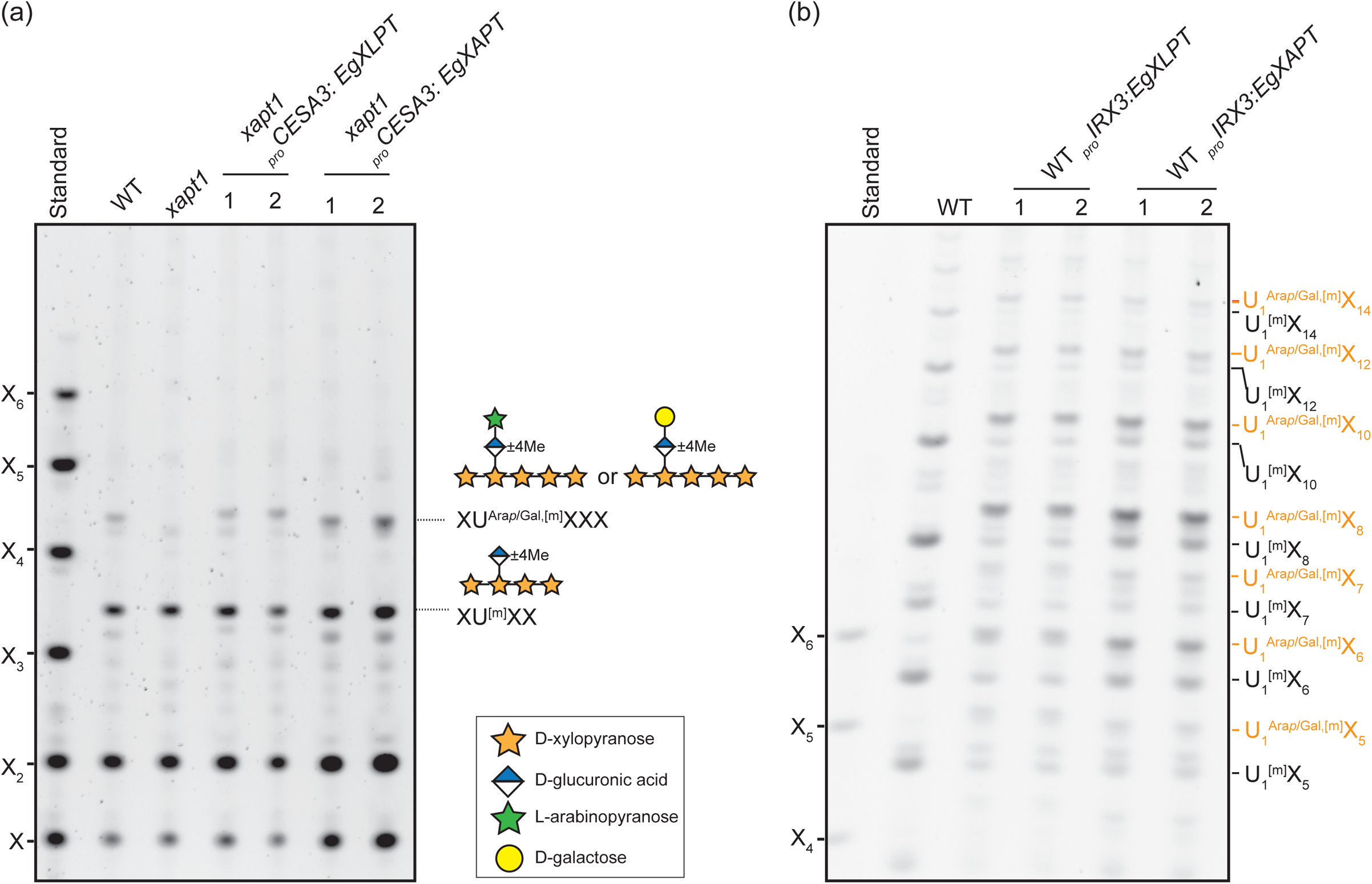
Overexpression of *Eg*XLPT or *Eg*XAPT can decorate the glucuronoxylan with an additional sugar. (a), Expression of *Eg*XLPT and *Eg*XAPT under primary cell wall *CESA3* promoter can complement the xapt1 xylan structure phenotype. Alkali-extracted xylan from 5-week-old young stem was digested with *Np*Xyn11A xylananase and analyzed by PACE. (b), Expression of *Eg*XLPT and *Eg*XAPT under secondary wall *IRX3* promoter in Arabidopsis WT can decorate most of the glucuronoxylan with an addition sugar. Alkali-extracted xylan from 8-week-old bottom stem was digested with *Bo*XynC xylanase and analysed by PACE. Xylo-oligosaccharide standard: X–X_6_.

PCW xylan is the presumed natural substrate of XAPT in Arabidopsis, but PCW xylan scarcity *in planta* makes its structure difficult to analyse. For this reason, we expressed the two *E. grandis* enzymes in SCW-producing tissues, which synthesise much more xylan. Furthermore, ^[Me]^GlcA sidechains on SCW xylan are completely unsubstituted (Bromley *et al*., 2013; Mortimer *et al*., 2015), which produces an ideal background for characterisation of *Eg*XAPT and *Eg*XLPT. The *Eg*XAPT and *Eg*XLPT coding sequences were placed under the control of the SCW-synthesis-specific *IRX3* promoter and expressed in WT Arabidopsis, and xylan from mature bottom-stems was digested with the glucuronoxylanase *Bo*XynC and analysed by PACE. As described previously (Bromley *et al*., 2013), digestion of the WT xylan released products decorated with unsubstituted GlcA (U_1_^[m]^X*_n_*), whose sizes corresponded to expected ^[Me]^GlcA spacings (Fig. 3b). In the *Eg*XAPT-and *Eg*XLPT-expressing lines, however, similar analysis revealed very different oligosaccharide products, suggesting that most of the ^[Me]^GlcA had been modified, perhaps into U^Ara*p*/Gal,[m]^ structures. The spacing between GlcA decorations along the xylan backbone did not appear to be affected in these transgenic plants. Interestingly, oligosaccharides with substituted GlcA in *Eg*XLPT-expressing lines migrated marginally slower than those in *Eg*XAPT-expressing lines, suggesting that the products of *Eg*XAPT and *Eg*XLPT might differ subtly in structure.

To further characterise the products of *Eg*XAPT and *Eg*XLPT, xylan from the relevant transgenic lines was digested simultaneously with *Np*Xyn11A and a *Bacteroides ovatus* GH115 (*Bo*GH115) α-glucuronidase, which removes unsubstituted GlcA side chains. As expected, PACE analysis indicated that WT bottom-stem xylan was fully digested into xylose and xylobiose (Fig. S3). In the *Eg*XAPT-and *Eg*XLPT-expressing lines, however, a band of higher molecular weight (XU^Ara*p*,[m]^XXX/XU^Gal,[m]^XXX) was produced. To distinguish between any arabinopyranosylated and galactosylated oligosaccharides, we then treated these products with a β-galactosidase (*An*GH35). Whereas the higher-molecular-weight band from *Eg*XAPT-expressing lines was completely insensitive to *An*GH35 (revealing it to be entirely XU^Ara*p*,[m]^XXX), almost all of the equivalent product from *Eg*XLPT-expressing lines shifted to XU^[m]^XXX after β-galactosidase treatment—revealing its identity as XU^Gal,[m]^XXX and indicating the presence of the U^Gal^ structure in these plants. These results suggest that *Eg*XLPT, but not *Eg*XAPT, is capable of transferring galactose to GlcA side chains.

To confirm these assignments, we analysed *Bo*XynC glucuronoxylanase products by mass spectrometry (MS), which can distinguish between pentose and hexose substituents. To prevent further ambiguity between U_1_^[m]^X*_n_* and U_1_^Ara*p*,[m]^X*_n_*_−1_, *Bo*XynC products were treated with α-glucuronidase *Bo*GH115 to remove unsubstituted GlcA residues, thereby leaving U_1_^Ara*p*/Gal^X*_n_* oligosaccharides as the only acidic products. The resulting oligosaccharides were derivatised with 2-aminobenzoic acid (2-AA) and then analysed by MALDI-TOF MS (Fig. 4, S4). As predicted, only xylo-oligosaccharides were produced from WT, confirming full removal of unsubstituted ^[Me]^GlcA by the α-glucuronidase. Products from plants expressing *At*XAPT under the same promoter were analysed in parallel for comparison. Lines expressing *At*XAPT and *Eg*XAPT exhibited abundant amounts of U_1_^Ara*p*^X*_n_* and low amounts of U_1_^Ara*p*,m^X*_n_*, confirming that *Eg*XAPT is a xylan arabinopyranosyltransferase. In lines expressing *Eg*XLPT, however, the dominant sugars released could be assigned as U_1_^Gal^X*_n_*, with a smaller presence of U_1_^Gal,m^X*_n_*. Interestingly, this approach also detected low amounts of U_1_^Ara*p*^X*_n_*and U_1_^Ara*p*,m^X*_n_* in all *Eg*XLPT independent lines (Fig. 4, S4). These results suggest that *Eg*XLPT is primarily a xylan galactosyltransferase with a low level of arabinopyranosyltransferase activity, at least when expressed in Arabidopsis.

**Figure 4.**
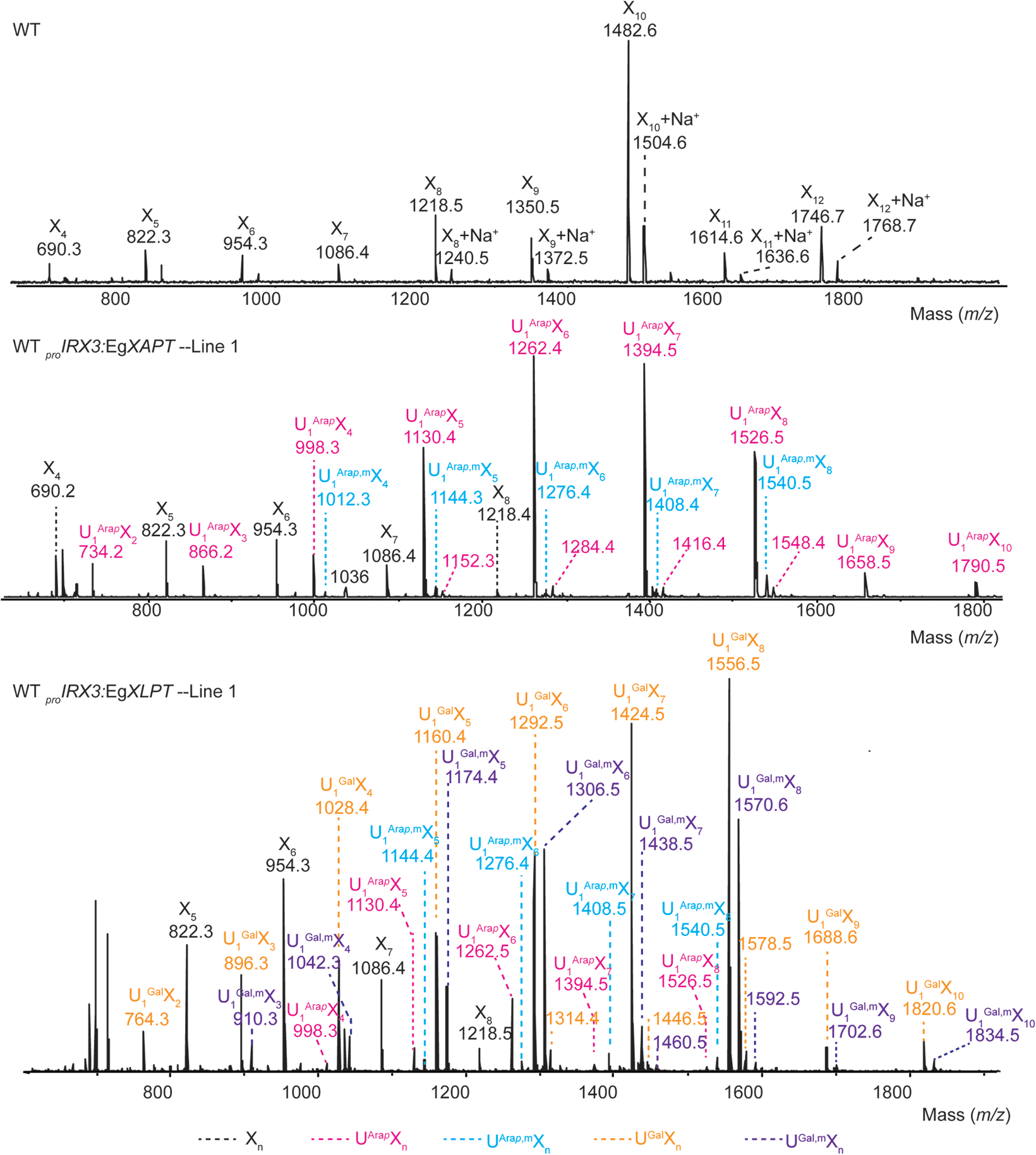
*Eg*XAPT and *Eg*XLPT exhibit different donor sugar specificities. AIR was extracted from 8-week-old bottom stems of either Arabidopsis WT plants or transgenic plants expressing *Eg*XAPT or *Eg*XLPT underthe secondary cell wall-specific promoter of IRX3. Alkali-extracted xylan was then digested with *Bo*Xyn30C xylanase followed by GH115 glucuronidase. The products were labelled with 2-AA and analysed by MALDI-TOF MS.

### Different *Eucalyptus dalrympleana* tissues contain structurally distinct xylans

Although U^Ara*p*^ and U^Gal^ side chains are known to exist in plants from the *Eucalyptus* genus (Togashi *et al*., 2009; Pena *et al*., 2016), the tissue distribution of these structures is unknown, limiting our insight into their function. To investigate where the enzymes might be active, we studied expression data available for *EgXAPT* and *EgXLPT* at https://EucGenIE.org (Hefer *et al*., 2011), which provides mRNA-Seq datasets for both *E. grandis* and the hybrid *E. grandis* × *E. urophylla*. Despite some potential disagreement between the two datasets, a clear pattern was still apparent: whereas *EgXAPT* expression is relatively constant and lowest in xylem tissues, the expression of *EgXLPT* is much stronger in xylem tissues than in leaves, flowers, and shoot tips (Fig. S5, S6). Interestingly, the expression of *EgXLPT* is dramatically higher in *E. grandis* × *E. urophylla* tension wood xylem than control xylem, as has been previously noted (Mizrachi *et al*., 2015).

To study the predicted xylan structure distributions *in planta*, we harvested an actively growing branch of *Eucalyptus dalrympleana* (mountain gum). To confirm that the *XAPT* and *XLPT* genes of *E. dalrympleana* do not differ substantially to those of *E. grandis*, we PCR-amplified them from genomic DNA using homology-based primers and sequenced the two intronless coding sequences, revealing 98% (XAPT) and 99% (XLPT) protein identity between the *E. dalrympleana* and *E. grandis* sequences (Fig. S7, S8). Next, to determine the extent of GlcA substitution amongst *E. dalrympleana* tissues, we sectioned the branch into cortex, phloem, sapwood/xylem, and leaf. Major leaf veins were removed to avoid contamination from vascular tissue (Fig. 5a). Alkali-treated AIR from each tissue was digested with *Bo*XynC xylanase and analysed with PACE. Based on the PACE xylo-oligosaccharide profiles, the tissues broadly fell into two groups with very different xylan structures (Fig. 5b). In the first group, which comprises leaf and phloem, U_1_^[m]^X_6_ and U_1_^Ara*p*/Gal,[m]^X_6_ were the most abundant xylan oligosaccharides, with traces of U_1_ ^[m]^X_5_ and U_1_^Ara*p*/Gal,[m]^X_5_. In the other group, which comprises cortex and xylem, the oligosaccharides were dominated by xylo-oligosaccharides decorated with single ^[Me]^GlcA. The cortex and xylem xylan had a GlcA spacing pattern very similar to SCW xylan from Arabidopsis (Bromley *et al*., 2013). Small amounts of U_1_^Ara*p*/Gal,[m]^X_5_ and U_1_^Ara*p*/Gal,[m]^X_6_ oligosaccharide could also be detected in cortex and xylem.

**Figure 5.**
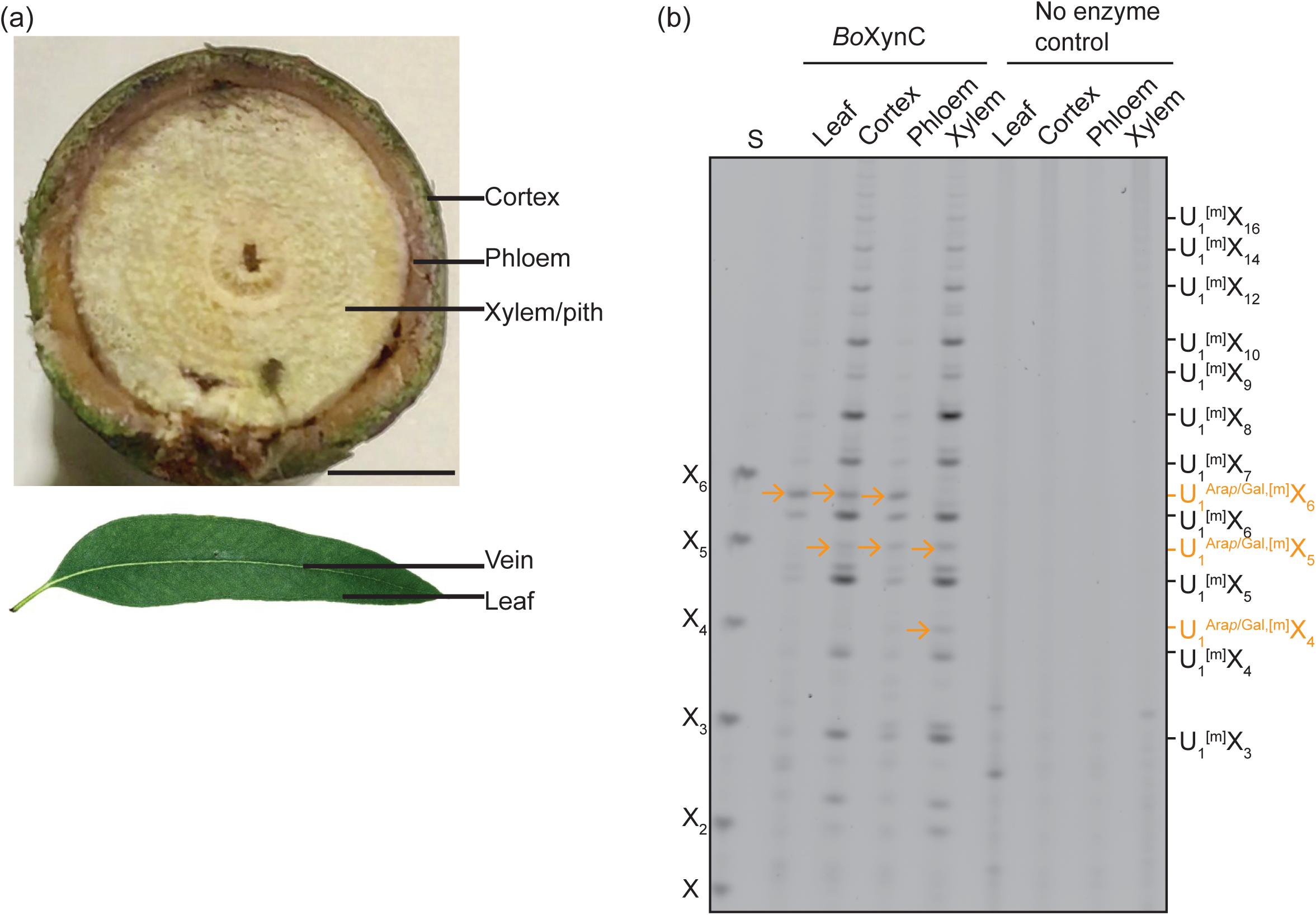
Xylan structure in different *Eucalyptus dalrympleana* tissues. (a), Eucalyptus stems were sectioned into cortex, phloem, and xylem/sapwood. Major leaf vein tissue was not included for the leaf xylan analysis. Scale bar = 0.9 cm. (b), PACE analysis of *Bo*XynC xylanase digests of AIR from different tissues. Xylo-oligosaccharide standard: X–X_6_. U^Ara*p*,Gal,[m]^X_n_ structures are labelled with arrows.

In order to discriminate U^Ara*p*^ and U^Gal^ structures in different Eucalyptus tissues, we again treated *Bo*XynC xylanase products with α-glucuronidase GH115 and analysed the remaining acidic products by MS. The MS profile of xylan showed that neutral and acidic oligosaccharides were released from all tissues but with different ratios (Fig. 6). Consistent with the expression data (Fig. S5, S6), leaf tissue released U_1_^Ara*p*,m^X_5_ and U_1_^Ara*p*,m^X_6_ oligosaccharides but no U_1_^Gal,m^X*_n_* oligosaccharides. In contrast, the stem tissues yielded detectable levels of U_1_^Gal,m^X_5_ and U ^Gal,m^X_6_. From cortex to phloem to xylem, the percentage of GlcA residues that were substituted decreased, while at the same time, the ratio of Gal to Ara*p* decorations increased. In summary, both the patterning and substitution of xylan GlcA side chains varies in different *E. dalrympleana* tissues and appears to correspond with the previously observed expression patterns of *Eg*XAPT and *Eg*XLPT in *E. grandis* / *E. grandis x E. urophylla*. More specifically, XAPT appears to be responsible for arabinosylating glucuronoxylan mainly in certain PCW-rich tissues, whereas glucuronoxylan galactosylation by XLPT appears to be enriched in woody tissues but is less abundant overall.

**Figure 6.**
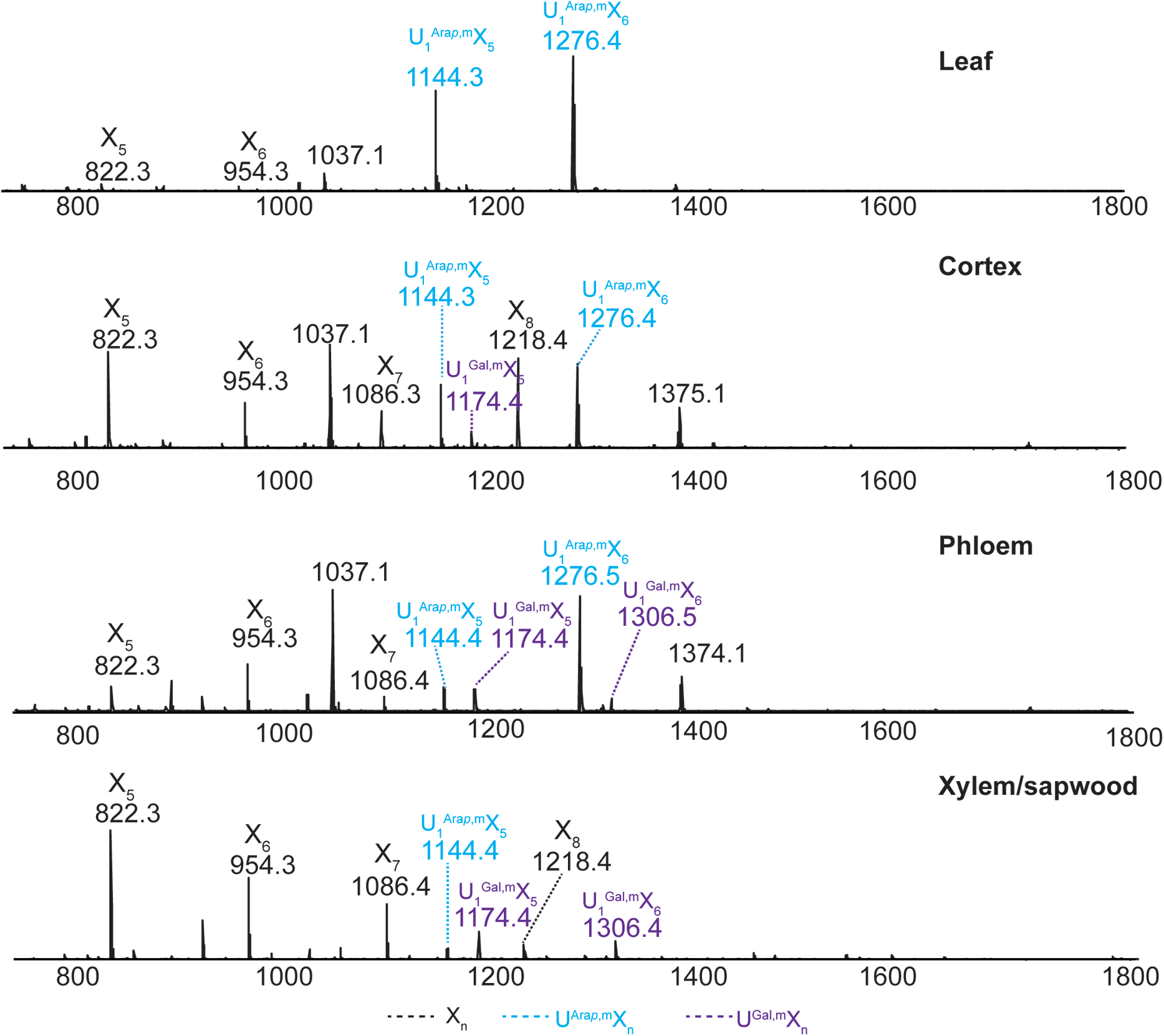
Distribution of xylan with decorated GlcA in different *Eucalyptus dalrympleana* tissues. AIR from different tissues was digested with *Bo*XynC xylanase followed by GH115 glucuronidase to remove the undecorated GlcA. The products were labelled with 2-AA and then analysed by MS.

### Other Myrtaceae plants contain U^Gal^ and U^Ara^*^p^* structures in their xylan

The results of our phylogenetic analysis suggested that other genera in the Myrtaceae family might possess XLPT orthologues, and therefore might exhibit U^Gal^ structures on their xylan. To investigate this hypothesis, we sectioned and analysed tissue from five additional Myrtaceae members: *Myrtus communis* (myrtle), *Metrosideros excelsa* (pohutukawa), *Plinia cauliflora* (jaboticaba), *Psidium guajava* (guava), and *Myrcianthes pungens* (guabiju). Xylan extracted from both cortex/phloem (combined) and xylem/pith was treated with xylanase *Np*Xyn11A and α-glucuronidase *Bo*GH115, and as before, the products were subsequently digested with *An*GH35 β-galactosidase. In xylem/pith, XU^Ara*p*,[m]^XXX/XU^Gal,[m]^XXX oligosaccharides were detectable in all species, with *M. communis* exhibiting remarkably high levels of GlcA substitution (Fig. S9). Notably, for *M. communis*, *P. cauliflora*, *P. guajava*, and *M. pungens* xylan, but not *M. excelsa*, a large proportion of this XU^Ara*p*,[m]^XXX/XU^Gal,[m]^XXX product was sensitive to *An*GH35, indicating that this band was primarily composed of the XU^Gal,[m]^XXX oligosaccharide with a smaller amount of XU^Ara*p*^XXX in xylem (Fig. S9). In cortex/phloem, XU^Ara*p*,[m]^XXX/XU^Gal,[m]^XXX oligosaccharides were detectable in all species except in *P. cauliflora* (Fig. S10). However, only *M. communis* yielded products that were sensitive to *An*GH35, suggesting a lack of U^Gal,[m]^ structures in the cortex/phloem of the other four species. Our results therefore support the idea that many Myrtaceae plants exhibit U^Gal^ side chains and that this structure is enriched in woodier tissues.

### GH30-family glucuronoxylanases have convergently evolved to overcome the obstacle of glucuronyl side chain modification

The distinct patterns of xylan arabinopyranosylation and galactosylation in Myrtaceae-family plants hint at functional importance for physical cell wall properties. We were therefore surprised to observe that Arabidopsis *xapt1* mutant plants exhibit no obvious growth phenotype; nor was any new phenotype revealed when *xapt1* plants were crossed with *mur3* plants lacking the xyloglucan β-galactosyltransferase MUR3 (Fig. S11). Nevertheless, we considered the possibility that xylan arabinopyranosylation could be important for other aspects of Arabidopsis biology. For example, cell wall polysaccharides constitute a barrier to potential pathogens and plant pathogens therefore rely on polysaccharide-degrading enzymes for their virulence. In this work, we have used two *endo*-xylanases to analyse xylan structure: *Np*Xyn11A, from CAZy family GH11, and *Bo*XynC, a member of GH30 subfamily 8 from the gut bacterium *B. ovatus*. In order to act on xylan, the latter of these specifically recognises and binds to GlcA substitutions. Curiously, previously solved crystal structures of related GH30 xylanases from plant pathogen *Dickeya chrysanthemi* (previously *Erwinia chrysanthemi*) strain D1 (*Ec*_D_Xyn30A) and *Bacillus subtilis* (*Bs*XynC) show that both enzymes make a hydrogen bond to the C2 hydroxyl of GlcA using the phenol group of Tyr_255_ or Tyr_231_ respectively (Fig. 7a) (St John *et al*., 2011; Urbanikova *et al*., 2011). We reasoned that the 1,2-linked Ara*p*/Gal substitution that decorates GlcA residues at this very hydroxyl should therefore impede *Ec*_D_Xyn30A and *Bs*XynC from binding their substrate. In spite of this, our results clearly showed that the related *Bo*XynC is able to bind and cleave at the U^Ara*p*/Gal^ decorations (Figs. 1b, 3b). To determine whether *Ec*_D_Xyn30A and other GH30 xylanases might have the same activity, we digested xylan from Arabidopsis *gux1,2* young stem with *Ec*_D_Xyn30A or *Ct*Xyn30A (from *Clostridium thermocellum*). PACE analysis of the products revealed that, whereas *Bo*XynC is capable of cleaving the xylan backbone adjacent to both substituted and unsubstituted ^[Me]^GlcA, *Ec*_D_Xyn30A and *Ct*Xyn30A are only able to cleave effectively at unsubstituted ^[Me]^GlcA decorations (Fig. 8a). These results demonstrate a hitherto unknown substrate specificity variant amongst GH30 glucuronoxylanases.

**Figure 7.**
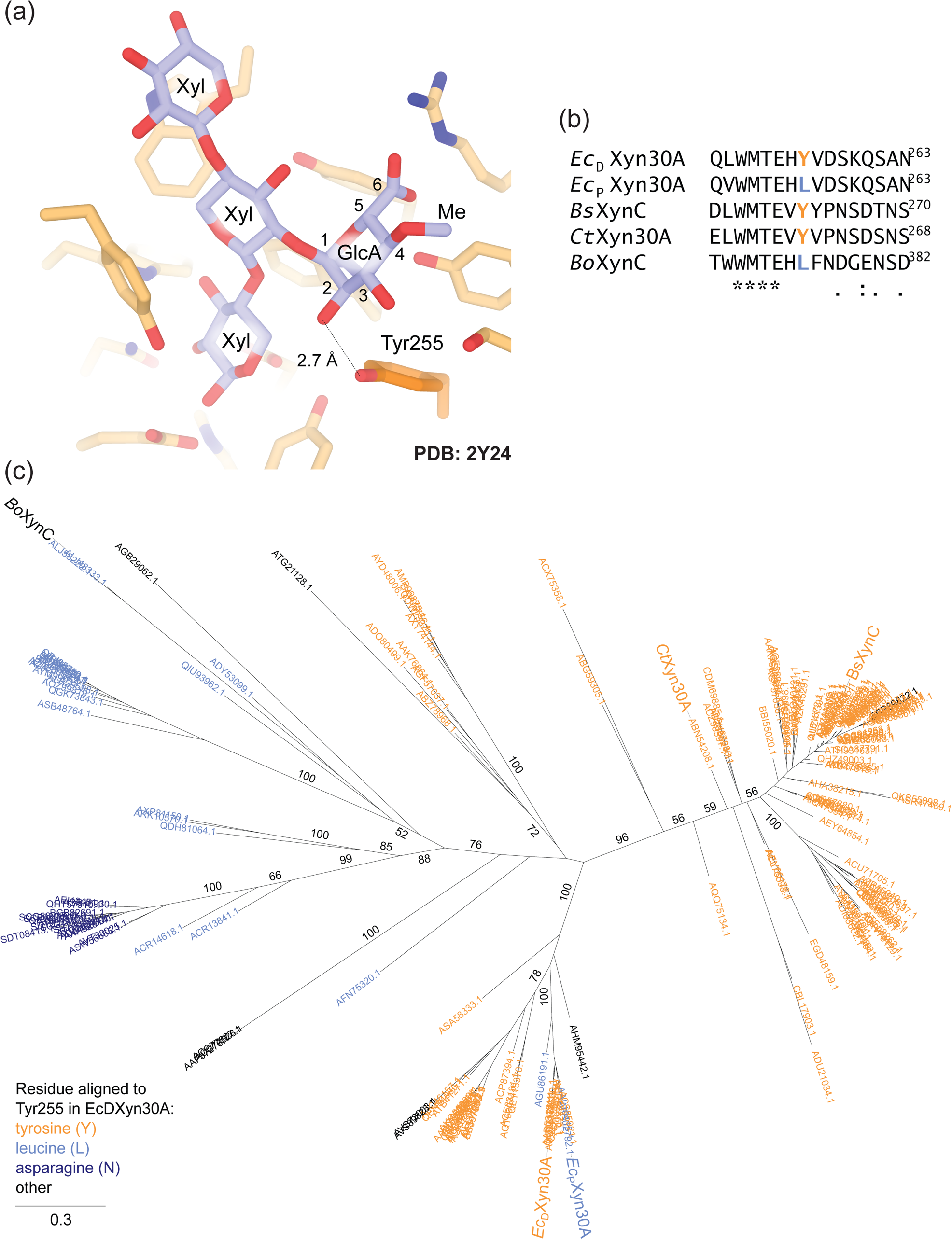
Differences in GH30 xylanase active site residues may reflect differences in ability to accommodate substituted glucuronic acid sidechains. (a), Previously characterised active site of GH30 xylanase from plant pathogen *Dickeya chrysanthemi* (previously *Erwinia chrysanthemi*) strain D1 (*Ec*_D_Xyn30A; PDB: 2Y24). Bound aldotetraouronic acid is shown in stick representation. The side chain of Tyr255, which forms a hydrogen bond with the GlcA C2 hydroxyl, is highlighted in orange. (b), Alignment segment of five GH30 xylanases, highlighting residues aligned to Tyr255 in *Ec*_D_Xyn30A. (c), Molecular phylogeny of GH30 subfamily 8 sequences downloaded from dbCAN2 (http://bcb.unl.edu/dbCAN2/). Sequences that display a tyrosine residue in the column aligned to Tyr255 are highlighted in orange; those that instead display leucine are highlighted in blue, while those with asparagine are highlighted in purple.

**Figure 8.**
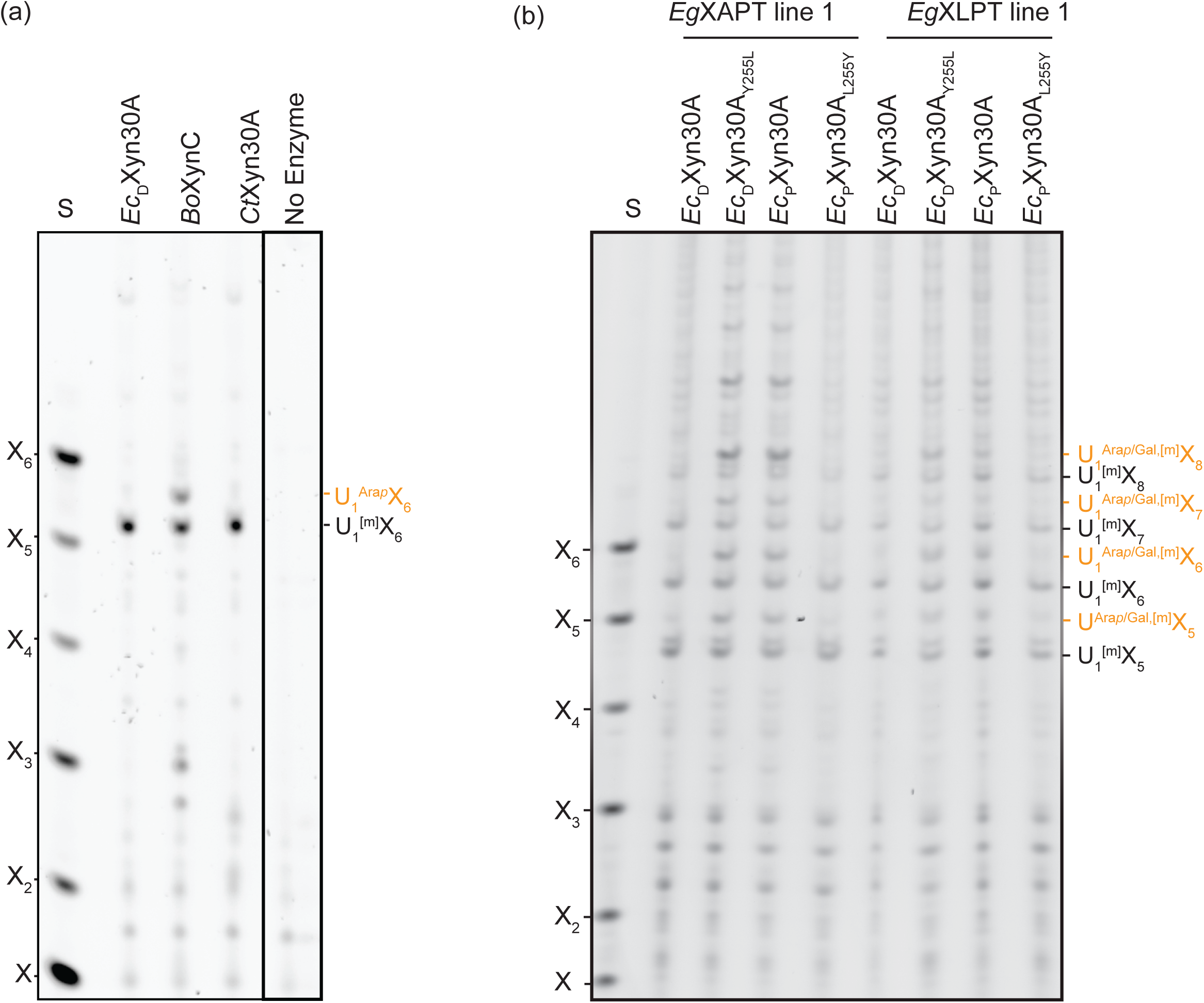
PACE gels to show the activities of different GH30 xylanases towards substituted glucuronoxylan. (a), PACE analysis of three GH30 xylanases-digests of xylan from *gux1 gux2* double mutant, which only contains primary cell wall glucuronoxylan. (b), AIR from the transgenic plants expressing *Eg*XAPT or *Eg*XLPT under the secondary cell wall-specific promoter of *IRX3* were used as the substrates. Alkali-extracted xylan was then digested with different GH30 and the corresponding modified GH30 enzymes. Xylo-oligosaccharide standard: X–X_6_.

To help understand the difference in substrate specificity between these enzymes, we aligned their protein sequences. The alignment showed that the tyrosine residue involved in binding the C2 hydroxyl of GlcA is also present in *Ct*Xyn30A. Interestingly, however, the corresponding residue in *Bo*XynC is not tyrosine but leucine (Fig. 7b).

To investigate the extent of amino acid variety at this position, we conducted a phylogenetic analysis of the entire GH30 subfamily 8. We then labelled the sequences in the tree according to the amino acid at the position of interest. The results showed that *Bo*XynC belongs to a distinct subgroup of enzymes with leucine rather than tyrosine at this position (Fig. 7c). Interestingly, we also identified an enzyme from *D. chrysanthemi* P860219 that diverged much more recently from *Ec*_D_Xyn30A than *Bo*XynC, but has independently acquired a leucine at this position (we henceforth refer to this enzyme as *Ec*_P_Xyn30A) (Fig. 7b, c). Although *Ec*_D_Xyn30A and *Ec*_P_Xyn30A otherwise share 82% protein sequence identity, like *Bo*XynC, *Ec*_P_Xyn30A (but not *Ec*_D_Xyn30A) was able to cut xylan adjacent to U^Ara*p*/Gal^ decorations (Fig. 8b).

To investigate whether this single residue difference is sufficient to explain the discrepancy in substrate specificity, we performed site-directed mutagenesis on the *Ec*_D_Xyn30A and *Ec*_P_Xyn30A enzymes. Leu_255_ from *Ec*_P_Xyn30A was mutated to tyrosine to produce *Ec*_P_Xyn30A_L255Y_, while Tyr_255_ from *Ec*_D_Xyn30A was mutated to leucine to produce *Ec*_D_Xyn30A_Y255L_. Initially, we used the purified enzymes to digest WT bottom-stem xylan without Ara*p*/Gal substitutions. All four enzymes released identical products of equal abundance, suggesting that all four variants exhibit glucuronoxylanase activity (Fig. S12). We used the purified enzymes to hydrolyse alkali-extracted xylan from plants expressing the *_pro_IRX3*:*EgXLPT*/*XAPT* constructs. As predicted, both *Ec*_D_Xyn30A and *Ec*_P_Xyn30A_L255Y_ produced primarily xylo-oligosaccharides with GlcA residues, whereas the enzymes *Ec*_D_Xyn30A_Y255L_ and *Ec*_P_Xyn30A could release many more products with U^Ara*p*/Gal^ decorations (Fig. 8b). These data suggest that a tyrosine residue at this position prevents efficient activity on xylan substrates with substituted GlcA decorations (or that leucine is required at this position for activity), and that multiple GH30 glucuronoxylanase subfamilies have independently adapted to accommodate substituted GlcA decorations. Importantly, these mutations appear to constitute counter-adaptations to the evolution of xylan arabinopyranosylation, implicating U^Ara*p*^ side chains in passive defence against pathogens.

## Discussion

In this work, we successfully predicted and characterised the enzymes responsible for decorating GlcA xylan branches in Arabidopsis and *Eucalyptus* spp., identifying GT47-A_V_ as a novel gene subfamily. While the two new activities share the most characteristic feature of others in GT47-A—adding a 2-linked sugar to an α-linked decoration of a hemicellulose backbone—it is notable that both donor (arabinopyranose) and acceptor (xylan) contrast with those of their closest characterised sisters (galactose/xyloglucan for both GT47-A_III_ and GT47-A_VI_) (Yu *et al*., 2022). This significant change in activity over a comparatively short timespan contributes to an emerging picture of rapid neofunctionalisation within the GT47-A subfamily (Yu *et al*., 2022; Wilson *et al*., 2023), whose driving forces we are just beginning to understand.

The identity of the pentosyl residue decorating GlcA branches in Arabidopsis PCW xylan has not been fully confirmed: previous attempts to identify it by NMR could not distinguish between α1,2-linked arabinopyranose and β1,2-linked xylose. Nevertheless, the former is favoured because the decoration is insensitive to β-xylosidase (Mortimer *et al*., 2015). Furthermore, the equivalent pentosyl decorations in *Eucalyptus grandis* and non-commelinid monocots have been unambiguously determined to be α1,2-linked arabinopyranose (Pena *et al*., 2016). The close evolutionary relationship between the Arabidopsis and *E. grandis* XAPT enzymes further strengthens the idea that the Arabidopsis pentosyl decoration is indeed arabinopyranose. Hence, we are confident in assigning *At*XAPT1 as an arabinopyranosyltransferase.

The identification of XAPT and XLPT enzymes in GT47-A_V_ helps to explain the distribution of substituted GlcA xylan side chains amongst plant species. For instance, our previous phylogeny (Yu *et al*., 2022) demonstrated that GT47-A_V_ genes have been completely lost from grass (Poaceae) genomes, explaining why U^Ara*p*^ side chains are present in non-commelinid monocot orders but have never been seen in grasses (we can also now predict the presence of U^Ara*p*^ structures in non-grass commelinids such as pineapple). We also observed candidate XAPT orthologues in lycophytes and gymnosperms; it will be interesting to determine if these species also carry decorated GlcA side chains. In addition, our work highlights the emergence of U^Gal^ side chains in the Myrtaceae family, likely arising from recent duplication and neofunctionalisation of the *XAPT* locus during the evolution of the Myrtales order.

Unlike arabinoxylan, glucuronoxylan is present in virtually every land plant cell wall, suggesting a specific and important role for GlcA in cell wall structure—perhaps as a covalent bridge to lignin polymers in secondary cell walls (Giummarella *et al*., 2019; Terrett & Dupree, 2019) or as a factor in xylan solubility and assembly onto cellulose (Schultink *et al*., 2015). Further modification of GlcA by 2-*O*-arabinopyranosylation, 2-*O*-galactosylation, and/or 4-*O*-methylation could therefore potentially modulate xylan incorporation into the cell wall. At the same time, the U^Ara*p*^ and U^Gal^ disaccharide side chains resemble the functionally important side chains of other PCW hemicelluloses: namely, the Gal-β1,2-Xyl-α1,6-and Gal-β1,2-Gal-α1,6-side chains of xyloglucan and β-GGM, respectively. For xyloglucan at least, the galactosyl decorations may influence its binding to cellulose (Zhao *et al*., 2014) and/or increase its solubility during biosynthesis (Tamura *et al*., 2005; Kong *et al*., 2015). Whether decoration of GlcA could have an effect on xylan–cellulose interactions will require extensive computational and solid state NMR work to establish.

Despite these possibilities, but consistent with GlcA methyltransferase mutants (Urbanowicz *et al*., 2012; Li *et al*., 2013) and glucuronyltransferase mutants (Mortimer *et al*., 2015), we did not observe any obvious phenotype of *xapt* or *xapt mur3-3* plants. Hence, any critical role for U^Ara*p*^ structures in Arabidopsis vegetative growth is unclear. Nevertheless, we observed an interesting interplay between GlcA arabinopyranosylation/galactosylation and methylation: whereas all U^Ara*p*^ and U^Gal^ structures in *E. dalrympleana* were methylated (i.e. U^Ara*p*,m^ and U^Gal,m^), the equivalent structures in transgenic Arabidopsis lines expressing *Eg*XAPT and *Eg*XLPT were almost completely unmethylated. This might suggest that Arabidopsis GlcA 4-*O*-methyltransferases do not accept substituted GlcA residues. Furthermore, some clues for the function of U^Ara*p*^ and U^Gal^ structures may lie in their uneven tissue distribution. Based on previous comparisons between ‘young’ and ‘old’ stems in Arabidopsis, U^Ara*p*^ structures have so far been assumed to be a general feature of ‘PCW xylan’ (Mortimer *et al*., 2015). However, in the Myrtaceae family at least, we have shown that the situation is likely more complicated. For example, while U^Ara*p*^ structures could be found in typically PCW-rich organs and tissues such as leaf, cortex, and phloem, U^Gal^ structures were specifically enriched in xylem/pith but completely absent from leaves. Furthermore, given that our dissections lacked the precision to separate individual cell types, it is possible that substitution of GlcA is limited to specialised cells within the tissues of interest. Single-cell expression atlases for Arabidopsis stem (Shi *et al*., 2021) and leaf (Kim *et al*., 2021) lend support to this idea by suggesting that *At*XAPT expression may be selectively enriched in cell types such as cambium, phloem parenchyma, and/or stem epidermis. Moreover, although we did not investigate the xylan structures *in planta*, *Eg*XLPT appears to be specifically upregulated in tension wood xylem tissues—perhaps by as much as a 50-fold increase (Mizrachi *et al*., 2015)—hinting at a specific functional role in the cell wall. As previously proposed, it is possible that xylan structure could be remodelled in tension wood, perhaps in a way that affects its incorporation into the wall. Whether any significant biochemical differences exist between U^Ara*p*^ and U^Gal^ structures is unclear, but it is possible that they differ in sensitivity to *exo*-glycosidases—be they microbial or cell wall remodelling enzymes (such as the β-galactosidase MUM2).

Whether or not the GlcA branches on xylan play a structural role in the cell wall, their absence leads to a major reduction in biomass resistance to enzymatic saccharification (Lyczakowski *et al*., 2017), suggesting that they may constitute a direct or indirect factor in plant-pathogen interaction. Here, we have uncovered that the substitution of GlcA residues likely affects the ability of GH30-family glucuronoxylanases to degrade xylan. Remarkably, different lineages of this family appear to have convergently overcome the obstacle posed by C2 modification through a single point mutation. Interestingly, the strain of *D. chrysanthemi* whose glucuronoxylanase has recently acquired tolerance for these decorations, P860219, was apparently cultured from *Anthurium* sp.: a non-commelinid from the Alismatales order whose close relatives exhibit abundant U^Ara*p*^ xylan side chains (Hurlbert & Preston, 2001). This is in contrast to substitution-intolerant strain D1, which was cultured from corn (Vroemen *et al*., 1995), a member of the Poaceae that lacks any substitutions. This hints that cell wall structures could be applying a direct selection pressure on xylanase evolution.

In summary, we have identified one of the missing components of xylan synthesis in higher plants. The identification of XAPT1 permits engineering of existing xylan polysaccharides using synthetic biology approaches. Furthermore, by revealing previously unknown substrate specificities in GH30 glucuronoxylanases, our results could facilitate optimisation of industrial lignocellulosic biomass degradation. These results hint at the constant evolutionary battle between plants and pathogens.

## Acknowledgements and funding

We thank Paul Stearn (Jesus College, Cambridge), Ángela Cano and Sam Brockington (Cambridge University Botanic Garden), and Pedro Araújo (University of Campinas) for providing plant material.

This work was supported by the Leverhulme Trust Centre for Natural Material Innovation, the Biotechnology and Biological Sciences Research Council (BBSRC) of the UK as part of the OpenPlant Synthetic Biology Research Centre (Reference BB/L014130/1), The UKRI grant underwriting the ERC advanced grant (EVOCATE Function and evolution of plant cell wall architecture for sustainable technologies. EP/X027120/1), Innovate UK, the Cambridge BBSRC-DTP Programme (Reference BB/J014540/1), a Novo Nordisk Foundation grant Oxymist (Grant Number NNF20OC0059697), and a BBSRC iCASE studentship (Reference BB/M015432/1).

## Author contributions

LY conceived the original screen, designed the study, performed most polysaccharide analyses and plant transformations, and co-wrote the paper; LFLW conducted the phylogenetic analyses, contributed to the biochemical analysis, designed the xylanase point mutations, and co-wrote the paper; OMT carried out the molecular biology, conducted xylanase expression and purification, and co-wrote the paper; JWR identified differences in xylanase activities and supported the biochemical analysis and identification of Arabidopsis T-DNA insertion lines; JJL supported the biochemical analysis and molecular biology; XY made the double mutant; KBRMK contributed to the design and production of mutated xylanases; PD designed and supervised the study, interpreted data and co-wrote the paper.

## Data Availability

The data that support the findings of this study are available in the main text and in the Supporting Information for this article. Sequences for newly identified *Eucalyptus dalrympleana* XAPT and XLPT will be deposited in GenBank after acceptance. All plant material descried in this manuscript is available on a reasonable request from the corresponding authors.

## References

1. Altschul SF, Gish W, Miller W, Myers EW, Lipman DJ. 1990. Basic local alignment search tool. Journal of Molecular Biology 215(3): 403–410.

2. Anders N, Wilson LFL, Sorieul M, Nikolovski N, Dupree P. 2022. β-1, 4-Xylan backbone synthesis in higher plants: How complex can it be? Frontiers in Plant Science 13.

3. Bellincampi D, Cervone F, Lionetti V. 2014. Plant cell wall dynamics and wall-related susceptibility in plant–pathogen interactions. Frontiers in Plant Science 5: 228.

4. Beri D, York WS, Lynd LR, Pena MJ, Herring CD. 2020. Development of a thermophilic coculture for corn fiber conversion to ethanol. Nature Communications 11(1): 1937.

5. Bromley JR, Busse-Wicher M, Tryfona T, Mortimer JC, Zhang ZN, Brown DM, Dupree P. 2013. GUX1 and GUX2 glucuronyltransferases decorate distinct domains of glucuronoxylan with different substitution patterns. Plant Journal 74(3): 423–434.

6. Brown DM, Zhang Z, Stephens E, Dupree P, Turner SR. 2009. Characterization of IRX10 and IRX10-like reveals an essential role in glucuronoxylan biosynthesis in Arabidopsis. Plant Journal 57(4): 732–746.

7. Burton RA, Gidley MJ, Fincher GB. 2010. Heterogeneity in the chemistry, structure and function of plant cell walls. Nature Chemical Biology 6(10): 724–732.

8. Busse-Wicher M, Gomes TCF, Tryfona T, Nikolovski N, Stott K, Grantham NJ, Bolam DN, Skaf MS, Dupree P. 2014. The pattern of xylan acetylation suggests xylan may interact with cellulose microfibrils as a twofold helical screw in the secondary plant cell wall of Arabidopsis thaliana. Plant Journal 79(3): 492–506.

9. Carpenter EJ, Matasci N, Ayyampalayam S, Wu S, Sun J, Yu J, Jimenez Vieira FR, Bowler C, Dorrell RG, Gitzendanner MA, **et al.** 2019. Access to RNA-sequencing data from 1,173 plant species: The 1000 Plant transcriptomes initiative (1KP). Gigascience 8(10).

10. Chmelová D, Škulcová D, Ondrejovic M. 2019. Microbial xylanases and their inhibition by specific proteins in cereals. Kvasny Prumysl 65(4): 127–133-127-133.

11. Chong SL, Koutaniemi S, Juvonen M, Derba-Maceluch M, Mellerowicz EJ, Tenkanen M. 2015. Glucuronic acid in Arabidopsis thaliana xylans carries a novel pentose substituent. International Journal of Biological Macromolecules 79: 807-812.

12. Clough SJ, Bent AF. 1998. Floral dip: a simplified method for Agrobacterium-mediated transformation of Arabidopsis thaliana. Plant Journal 16(6): 735-743.

13. Darriba D, Taboada GL, Doallo R, Posada D. 2011. ProtTest 3: fast selection of best-fit models of protein evolution. Bioinformatics 27(8): 1164-1165.

14. Drula E, Garron ML, Dogan S, Lombard V, Henrissat B, Terrapon N. 2022. The carbohydrate-active enzyme database: functions and literature. Nucleic Acids Res 50(D1): D571-D577.

15. Eddy SR. 2011. Accelerated Profile HMM Searches. PLoS Comput Biol 7(10): e1002195.

16. Edgar RC. 2004a. MUSCLE: a multiple sequence alignment method with reduced time and space complexity. BMC Bioinformatics 5: 113.

17. Edgar RC. 2004b. MUSCLE: multiple sequence alignment with high accuracy and high throughput. Nucleic Acids Research 32(5): 1792-1797.

18. Evtuguin DV, Tomas JL, Silva AMS, Neto CP. 2003. Characterization of an acetylated heteroxylan from Eucalyptus globulus Labill. Carbohydrate Research 338(7): 597-604.

19. Fu L, Niu B, Zhu Z, Wu S, Li W. 2012. CD-HIT: accelerated for clustering the next-generation sequencing data. Bioinformatics 28(23): 3150-3152.

20. Giummarella N, Pu YQ, Ragauskas AJ, Lawoko M. 2019. A critical review on the analysis of lignin carbohydrate bonds. Green Chemistry 21(7): 1573-1595.

21. Goubet F, Barton CJ, Mortimer JC, Yu XL, Zhang ZN, Miles GP, Richens J, Liepman AH, Seffen K, Dupree P. 2009. Cell wall glucomannan in Arabidopsis is synthesised by CSLA glycosyltransferases, and influences the progression of embryogenesis. Plant Journal 60(3): 527-538.

22. Grantham NJ, Wurman-Rodrich J, Terrett OM, Lyczakowski JJ, Stott K, Iuga D, Simmons TJ, Durand-Tardif M, Brown SP, Dupree R, **et al.** 2017. An even pattern of xylan substitution is critical for interaction with cellulose in plant cell walls. Nature Plants 3(11): 859-865.

23. Han MH, Liu YT, Zhang FL, Sun DF, Jiang JX. 2020. Effect of galactose side-chain on the self-assembly of xyloglucan macromolecule. Carbohydrate Polymers 246.

24. Hefer C, Mizrachi E, Joubert F, Myburg A 2011. The Eucalyptus genome integrative explorer (EucGenIE): a resource for Eucalyptus genomics and transcriptomics. BMC Proceedings: Springer. 1-1.

25. Hurlbert JC, Preston JF, 3rd. 2001. Functional characterization of a novel xylanase from a corn strain of Erwinia chrysanthemi. Journal of Bacteriology 183(6): 2093-2100.

26. Immelmann R, Gawenda N, Ramirez V, Pauly M. 2023. Identification of a xyloglucan beta-xylopyranosyltransferase from Vaccinium corymbosum. Plant Direct 7(7): e514.

27. Izuno A, Hatakeyama M, Nishiyama T, Tamaki I, Shimizu-Inatsugi R, Sasaki R, Shimizu KK, Isagi Y. 2016. Genome sequencing of Metrosideros polymorpha (Myrtaceae), a dominant species in various habitats in the Hawaiian Islands with remarkable phenotypic variations. Journal of Plant Research 129(4): 727-736.

28. Izuno A, Wicker T, Hatakeyama M, Copetti D, Shimizu KK. 2019. Updated Genome Assembly and Annotation for Metrosideros polymorpha, an Emerging Model Tree Species of Ecological Divergence. G3 (Bethesda) 9(11): 3513-3520.

29. James CM, Barrett JA, Russell SJ, Gibby M. 2001. A rapid PCR based method to establish the potential for paternal inheritance of chloroplasts in Pelargonium. Plant Molecular Biology Reporter 19: 163-167.

30. Katoh K, Standley DM. 2013. MAFFT multiple sequence alignment software version 7: improvements in performance and usability. Molecular Biology and Evolution 30(4): 772-780.

31. Kim JY, Symeonidi E, Pang TY, Denyer T, Weidauer D, Bezrutczyk M, Miras M, Zollner N, Hartwig T, Wudick MM, **et al.** 2021. Distinct identities of leaf phloem cells revealed by single cell transcriptomics. Plant Cell 33(3): 511-530.

32. Kong YZ, Pena MJ, Renna L, Avci U, Pattathil S, Tuomivaara ST, Li XM, Reiter WD, Brandizzi F, Hahn MG, **et al.** 2015. Galactose-depleted xyloglucan is dysfunctional and leads to dwarfism in Arabidopsis. Plant Physiology 167(4): 1296-U1294.

33. Li W, Godzik A. 2006. Cd-hit: a fast program for clustering and comparing large sets of protein or nucleotide sequences. Bioinformatics 22(13): 1658-1659.

34. Li XF, Jackson P, Rubtsov DV, Faria-Blanc N, Mortimer JC, Turner SR, Krogh KB, Johansen KS, Dupree P. 2013. Development and application of a high throughput carbohydrate profiling technique for analyzing plant cell wall polysaccharides and carbohydrate active enzymes. Biotechnology for Biofuels 6:94.

35. Li XM, Cordero I, Caplan J, Molhoj M, Reiter WD. 2004. Molecular analysis of 10 coding regions from arabidopsis that are homologous to the MUR3 xyloglucan galactosyltransferase. Plant Physiology 134(3): 940-950.

36. Lyczakowski JJ, Wicher KB, Terrett OM, Faria-Blanc N, Yu X, Brown D, Krogh K, Dupree P, Busse-Wicher M. 2017. Removal of glucuronic acid from xylan is a strategy to improve the conversion of plant biomass to sugars for bioenergy. Biotechnology for Biofuels 10: 224.

37. Malinovsky FG, Fangel JU, Willats WG. 2014. The role of the cell wall in plant immunity. Frontiers in Plant Science 5: 178.

38. Mizrachi E, Maloney VJ, Silberbauer J, Hefer CA, Berger DK, Mansfield SD, Myburg AA. 2015. Investigating the molecular underpinnings underlying morphology and changes in carbon partitioning during tension wood formation in Eucalyptus. New Phytologist 206(4): 1351-1363.

39. Mortimer JC, Faria-Blanc N, Yu XL, Tryfona T, Sorieul M, Ng YZ, Zhang ZN, Stott K, Anders N, Dupree P. 2015. An unusual xylan in Arabidopsis primary cell walls is synthesised by GUX3, IRX9L, IRX10L and IRX14. Plant Journal 83(3): 413-426.

40. One Thousand Plant Transcriptomes I. 2019. One thousand plant transcriptomes and the phylogenomics of green plants. Nature 574(7780): 679-685.

41. Paes G, Berrin JG, Beaugrand J. 2012. GH11 xylanases: Structure/function/properties relationships and applications. Biotechnology Advances 30(3): 564-592.

42. Paradis E, Schliep K. 2019. ape 5.0: an environment for modern phylogenetics and evolutionary analyses in R. Bioinformatics 35(3): 526-528.

43. Patron NJ, Orzaez D, Marillonnet S, Warzecha H, Matthewman C, Youles M, Raitskin O, Leveau A, Farre G, Rogers C, **et al.** 2015. Standards for plant synthetic biology: a common syntax for exchange of DNA parts. New Phytologist 208(1): 13-19.

44. Pena MJ, Kulkarni AR, Backe J, Boyd M, O’Neill MA, York WS. 2016. Structural diversity of xylans in the cell walls of monocots. Planta 244(3): 589-606.

45. Pinto PC, Evtuguin DV, Neto CP. 2005. Structure of hardwood glucuronoxylans: modifications and impact on pulp retention during wood kraft pulping. Carbohydrate Polymers 60(4): 489-497.

46. Rogowski A, Briggs JA, Mortimer JC, Tryfona T, Terrapon N, Lowe EC, Basle A, Morland C, Day AM, Zheng HJ, **et al.** 2015. Glycan complexity dictates microbial resource allocation in the large intestine. Nature Communications 6: 7481.

47. Scheller HV, Ulvskov P. 2010. Hemicelluloses. Annual Review of Plant Biology 61: 263-289.

48. Schultink A, Cheng K, Park YB, Cosgrove DJ, Pauly M. 2013. The identification of two arabinosyltransferases from tomato reveals functional equivalency of xyloglucan side chain substituents. Plant Physiology 163(1): 86-94.

49. Schultink A, Naylor D, Dama M, Pauly M. 2015. The Role of the Plant-Specific ALTERED XYLOGLUCAN9 Protein in Arabidopsis Cell Wall Polysaccharide O-Acetylation. Plant Physiology 167(4): 1271-1283.

50. Shatalov AA, Evtuguin DV, Neto CP. 1999. (2-O-alpha-D-galactopyranosyl-4-O-methyl-alpha-D-glucurono)-D-xylan from Eucalyptus globulus Labill. Carbohydrate Research 320(1-2): 93-99.

51. Shi D, Jouannet V, Agusti J, Kaul V, Levitsky V, Sanchez P, Mironova VV, Greb T. 2021. Tissue-specific transcriptome profiling of the Arabidopsis inflorescence stem reveals local cellular signatures. Plant Cell 33(2): 200-223.

52. Simmons TJ, Mortimer JC, Bernardinelli OD, Pöppler AC, Brown SP, Deazevedo ER, Dupree R, Dupree P. 2016. Folding of xylan onto cellulose fibrils in plant cell walls revealed by solid-state NMR. Nature Communications 7: 13902.

53. St John FJ, Hurlbert JC, Rice JD, Preston JF, Pozharski E. 2011. Ligand bound structures of a glycosyl hydrolase family 30 glucuronoxylan xylanohydrolase. Journal of Molecular Biology 407(1): 92-109.

54. Stamatakis A. 2014. RAxML version 8: a tool for phylogenetic analysis and post-analysis of large phylogenies. Bioinformatics 30(9): 1312-1313.

55. Tamura K, Shimada T, Kondo M, Nishimura M, Hara-Nishimura I. 2005. KATAMARI1/MURUS3 is a novel Golgi membrane protein that is required for endomembrane organization in Arabidopsis. Plant Cell 17(6): 1764-1776.

56. Terrett OM, Dupree P. 2019. Covalent interactions between lignin and hemicelluloses in plant secondary cell walls. Current Opinion in Biotechnology 56: 97-104.

57. Togashi H, Kato A, Shimizu K. 2009. Enzymatically derived aldouronic acids from Eucalyptus globulus glucuronoxylan. Carbohydrate Polymers 78(2): 247-252.

58. Tryfona T, Bourdon M, Delgado Marques R, Busse-Wicher M, Vilaplana F, Stott K, Dupree P. 2023. Grass xylan structural variation suggests functional specialization and distinctive interaction with cellulose and lignin. Plant Journal 113(5): 1004-1020.

59. Tryfona T, Liang HC, Kotake T, Kaneko S, Marsh J, Ichinose H, Lovegrove A, Tsumuraya Y, Shewry PR, Stephens E, **et al.** 2010. Carbohydrate structural analysis of wheat flour arabinogalactan protein. Carbohydrate Research 345(18): 2648-2656.

60. Tryfona T, Stephens E. 2010. Analysis of carbohydrates on proteins by offline normal-phase liquid chromatography MALDI-TOF/TOF-MS/MS. Plant Cell Wall: Methods and Protocols 658: 137-151.

61. Urbanikova L, Vrsanska M, Morkeberg Krogh KB, Hoff T, Biely P. 2011. Structural basis for substrate recognition by Erwinia chrysanthemi GH30 glucuronoxylanase. FEBS Journal 278(12): 2105-2116.

62. Urbanowicz BR, Pena MJ, Ratnaparkhe S, Avci U, Backe J, Steet HF, Foston M, Li HJ, O’Neill MA, Ragauskas AJ, **et al.** 2012. 4-O-methylation of glucuronic acid in Arabidopsis glucuronoxylan is catalyzed by a domain of unknown function family 579 protein. Proceedings of the National Academy of Sciences of the United States of America 109(35): 14253-14258.

63. Van Bel M, Diels T, Vancaester E, Kreft L, Botzki A, Van de Peer Y, Coppens F, Vandepoele K. 2018. PLAZA 4.0: an integrative resource for functional, evolutionary and comparative plant genomics. Nucleic Acids Research 46(D1): D1190-D1196.

64. Vroemen S, Heldens J, Boyd C, Henrissat B, Keen NT. 1995. Cloning and characterization of the bgxA gene from Erwinia chrysanthemi D1 which encodes a beta-glucosidase/xylosidase enzyme. Molecular Genetics and Genomics 246(4): 465-477.

65. Wei QQ, Yang Y, Li H, Liu ZW, Fu R, Feng HQ, Li C. 2021. The xyloglucan galactosylation modulates the cell wall stability of pollen tube. Planta 254(6): 133-140.

66. Wilson LF, Neun S, Yu L, Tryfona T, Stott K, Hollfelder F, Dupree P. 2023. The biosynthesis, degradation, and function of cell wall β[xylosylated xyloglucan mirrors that of arabinoxyloglucan. New Phytologist 240 (6): 2353-2371.

67. Wu AM, Hornblad E, Voxeur A, Gerber L, Rihouey C, Lerouge P, Marchant A. 2010. Analysis of the Arabidopsis IRX9/IRX9-L and IRX14/IRX14-L pairs of glycosyltransferase genes reveals critical contributions to biosynthesis of the hemicellulose glucuronoxylan. Plant Physiology 153(2): 542-554.

68. Yu L, Yoshimi Y, Cresswell R, Wightman R, Lyczakowski JJ, Wilson LFL, Ishida K, Stott K, Yu XL, Charalambous S, **et al.** 2022. Eudicot primary cell wall glucomannan is related in synthesis, structure, and function to xyloglucan. Plant Cell 34(11): 4600-4622.

69. Zhang H, Yohe T, Huang L, Entwistle S, brown P, Yang Z, Busk PK, Xu Y, Yin Y. 2018. dbCAN2: a meta server for automated carbohydrate-active enzyme annotation. Nucleic Acids Research 46(W1): W95-W101.

70. Zhao Z, Crespi VH, Kubicki JD, Cosgrove DJ, Zhong LH. 2014. Molecular dynamics simulation study of xyloglucan adsorption on cellulose surfaces: effects of surface hydrophobicity and side-chain variation. Cellulose 21(2): 1025-1039.

71. Zhu L, Dama M, Pauly M. 2018. Identification of an arabinopyranosyltransferase from *Physcomitrella patens* involved in the synthesis of the hemicellulose xyloglucan. Plant Direct 2(3): e00046.

