## Supplemental Files for "Evolution of glucuronoxylan side chain variability in vascular plants and the counter-adaptation of pathogenic cell-wall-degrading hydrolases"

**Supporting information**

Article title: XAPT and XLPT enzymes modify the glucuronic acid side chains of tissue-specific xylans in Arabidopsis and Eucalyptus

Authors: Li Yu, Louis F.L. Wilson, Oliver M. Terrett, Joel Wurman-Rodrich, Jan J. Lyczakowski, Xiaolan Yu, Kristian B.R.M. Krogh, Paul Dupree

Article acceptance date: n/a

**The following supporting information is available for this article:**

### Fig. S1. Arap-GlcA- structure is lost in *xapt1* young stem.

**Fig. S2**. Full phylogenetic tree of XAPT clade.

**Fig. S3**. Galactosylated glucuronoxylan from *Eg*XLPT-expressing lines is sensitive to β-galactosidase.

**Fig. S4**. *Eg*XAPT and *Eg*XLPT show different enzyme specificities.

**Fig. S5**. Expression pattern of Eucgr.D00738, downloaded from <https://eucgenie.org/>.

**Fig. S6**. Expression pattern of Eucgr.H00343, downloaded from <https://eucgenie.org/>.

**Fig. S7**. *Ed*XAPT and *Eg*XAPT coding sequences show high similarity.

**Fig. S8**. *Ed*XLPT and *Eg*XLPT coding sequences show high similarity.

**Fig. S9**. Other Myrtaceae plants contain U^Gal,[m]^ and U^Ara^*^p^*^,[m]^ structures in their xylan.

**Fig. S10**. Other Myrtaceae plants contain U^Gal,[m]^ and U^Ara^*^p^*^,[m]^ structures in their xylan.

### Fig. S11. *xapt1* mutant plants show no obvious growth phenotype.

### Fig. S12. Xylan from WT bottom stem can be digested in the same way with *Ec*_D_Xyn30A, *Ec*_D_Xyn30A_Y255L_, *Ec*_P_Xyn30A, and *Ec*_P_Xyn30A_L255Y_.

**Table S1**. Primers used in this study.

### Supplementary Figures

### Fig. S1. Ara*p*-GlcA- structure is lost in *xapt1* young stem. Alkali-extracted xylan from five-week-old young stem AIR was digested with *Np*Xyn11A and analysed using PACE. The *irx9l* mutant was used as a control. Xylo-oligosaccharide standard: X–X_6_.

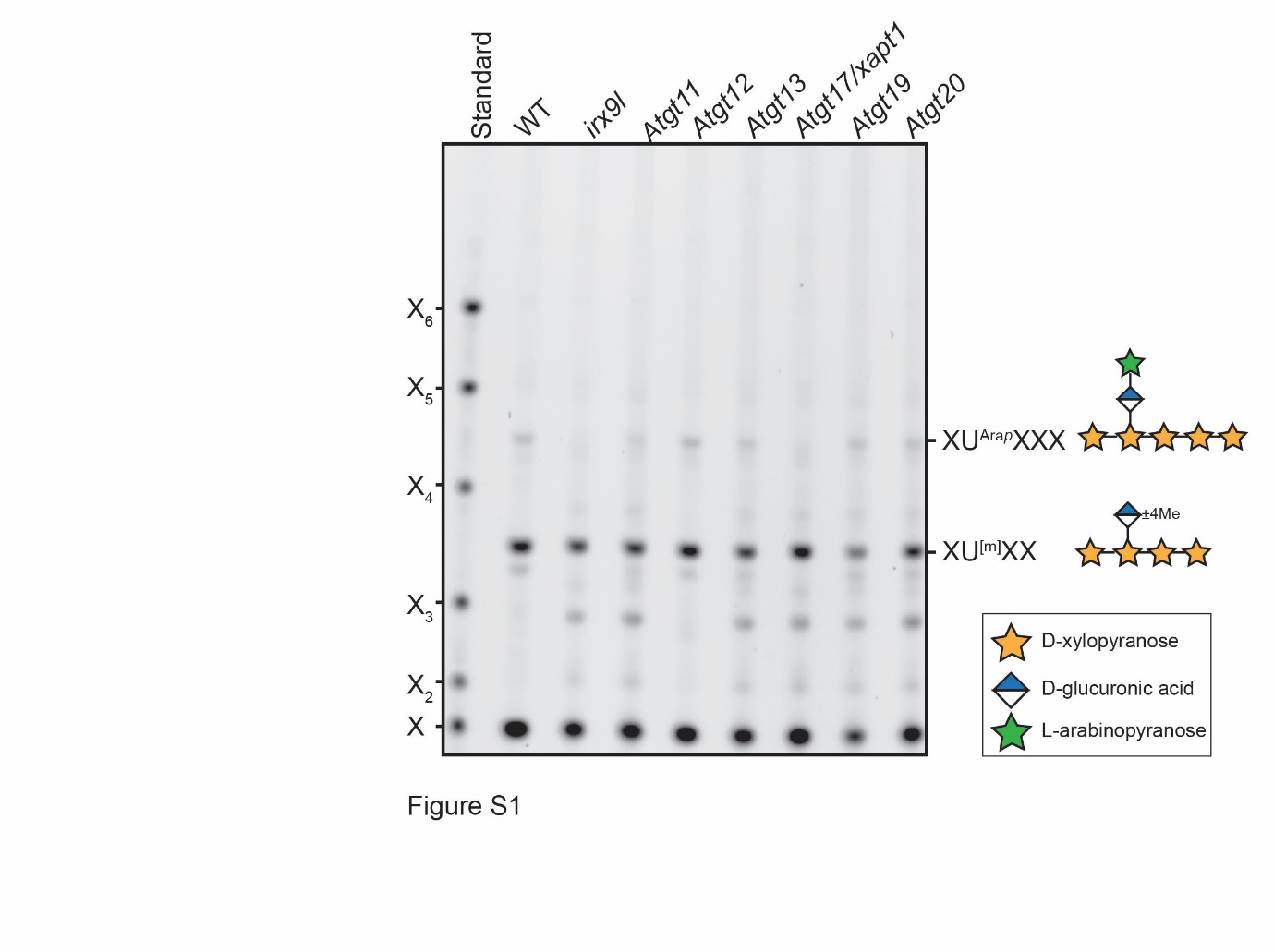

**Fig. S2. Full phylogenetic tree of XAPT clade.** See Figure 2 legend.

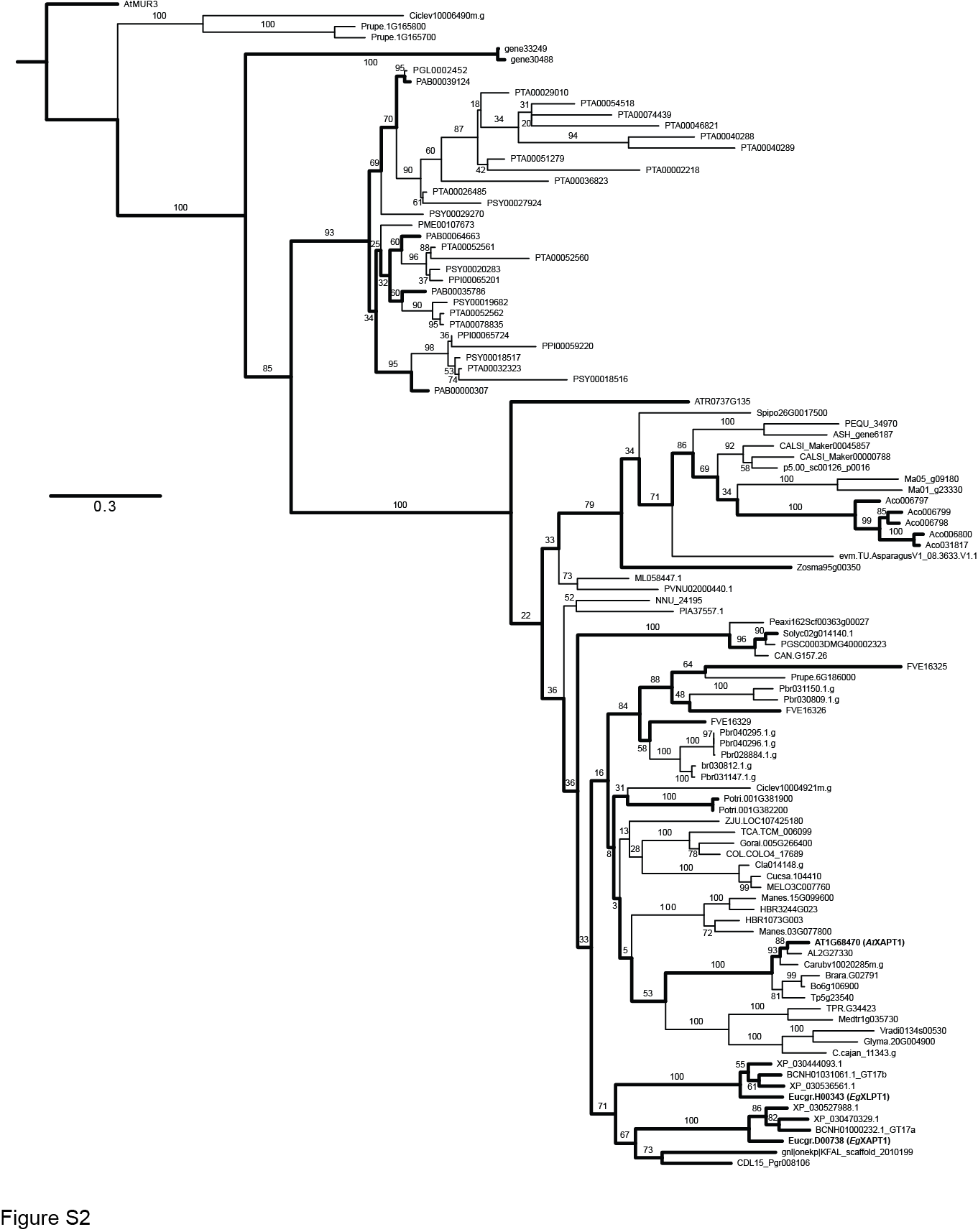

**Fig. S3. Galactosylated glucuronoxylan from *Eg*XLPT-expressing lines is sensitive to β-galactosidase.** Alkali-extracted xylan from Arabidopsis young stem AIR was digested simultaneously with *Np*Xyn11A xylanase and GH115 glucuronidase, which were subsequently removed by centrifugal filtration and ethanol precipitation. The resultant oligosaccharides were treated with or without *An*GH35 β-galactosidase. Xylo-oligosaccharide standard: X–X6.

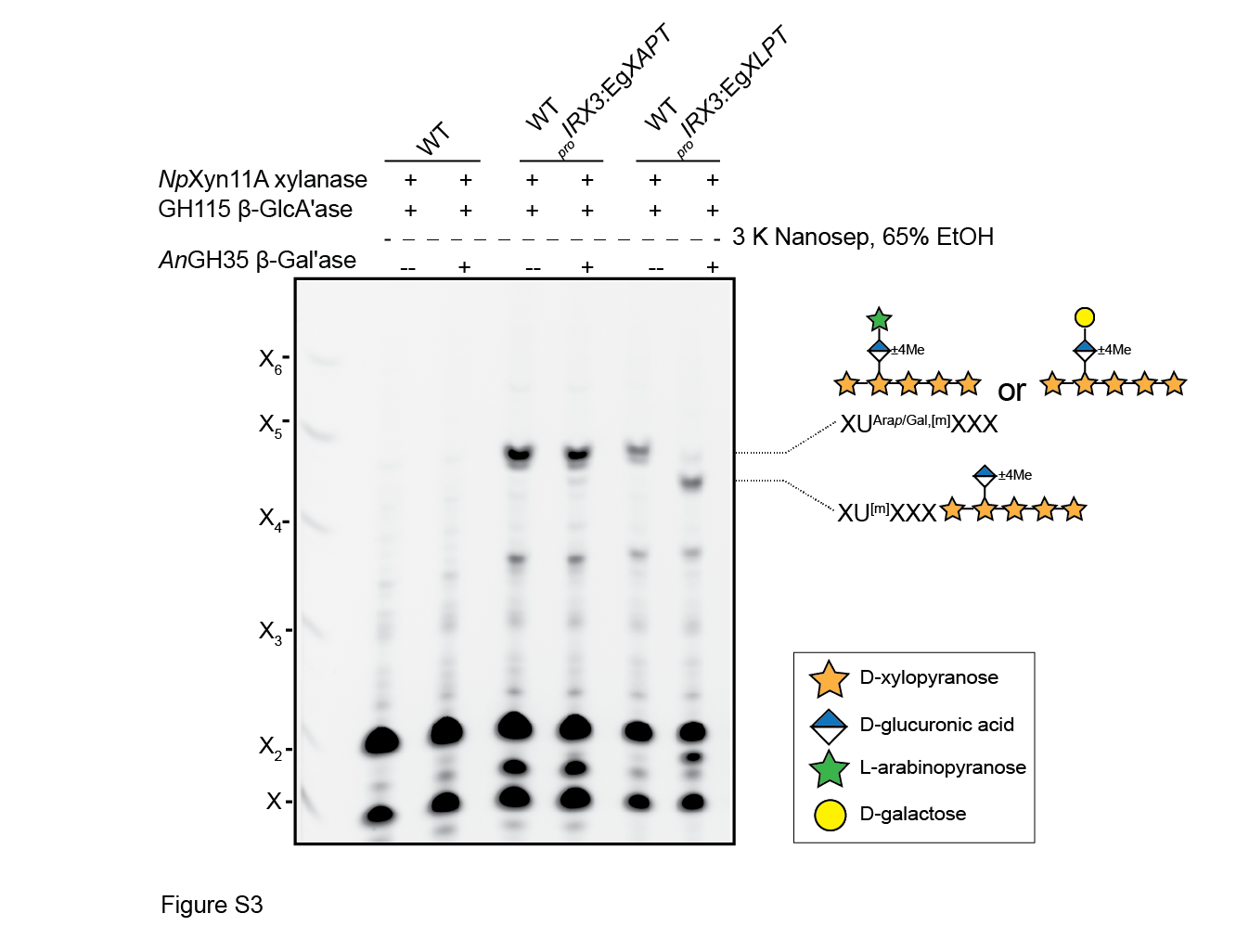

**Fig. S4. *Eg*XAPT and *Eg*XLPT show different enzyme specificities.** AIR was extracted from eight-week-old bottom stems of either WT plants or transgenic plants expressing *Eg*XAPT or *Eg*XLPT under the secondary cell wall-specific promoter of *IRX3*. Alkali-extracted xylan was then digested with *Bo*XynC glucuronoxylanase followed by GH115 glucuronidase. The products were labelled with 2-AA and analysed by MALDI-TOF MS.

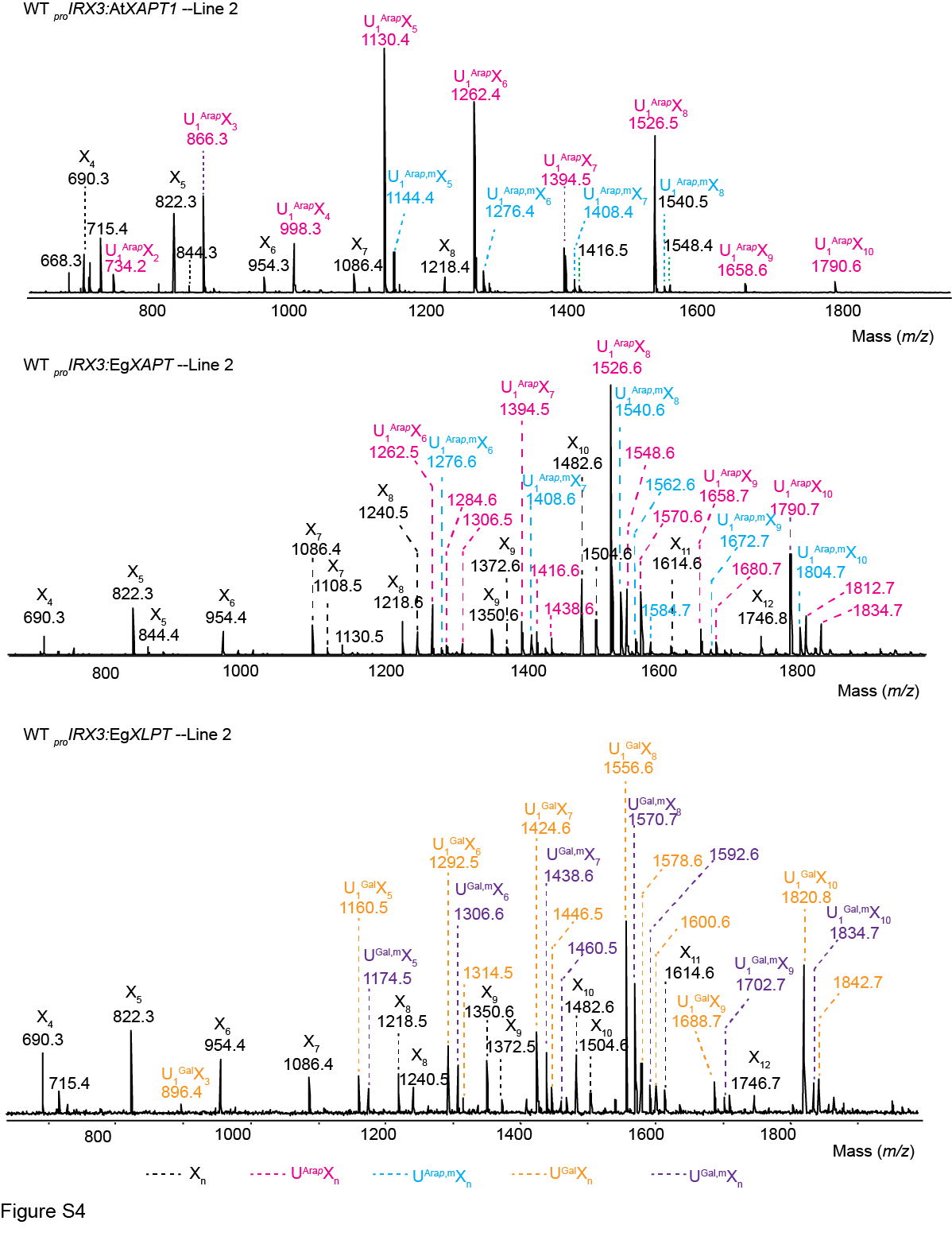

**Fig. S5. Expression pattern of Eucgr.D00738, downloaded from** [**https://eucgenie.org/**](https://eucgenie.org/)**.**

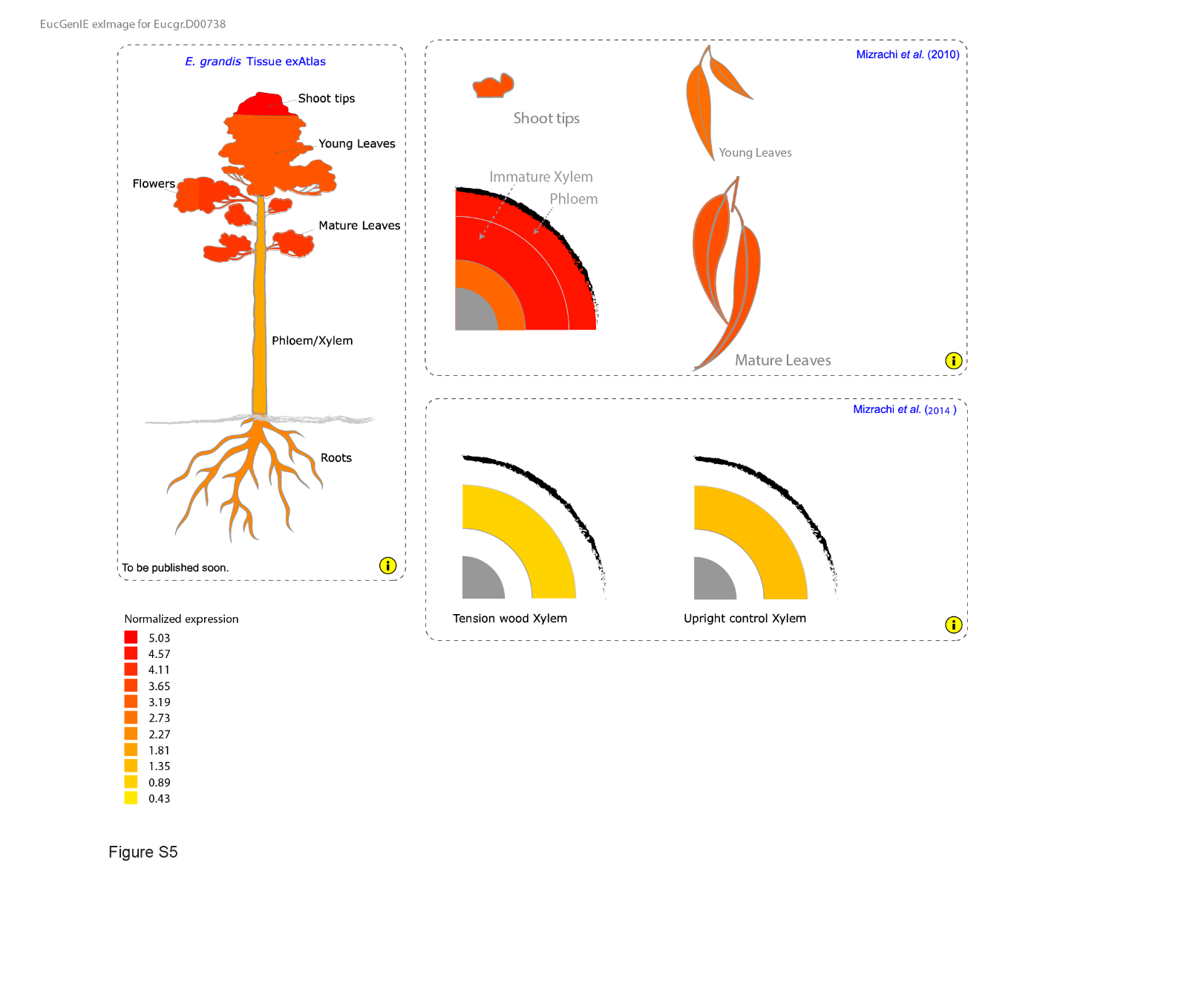

**Fig. S6. Expression pattern of Eucgr.H00343, downloaded from** [**https://eucgenie.org/**](https://eucgenie.org/)**.**

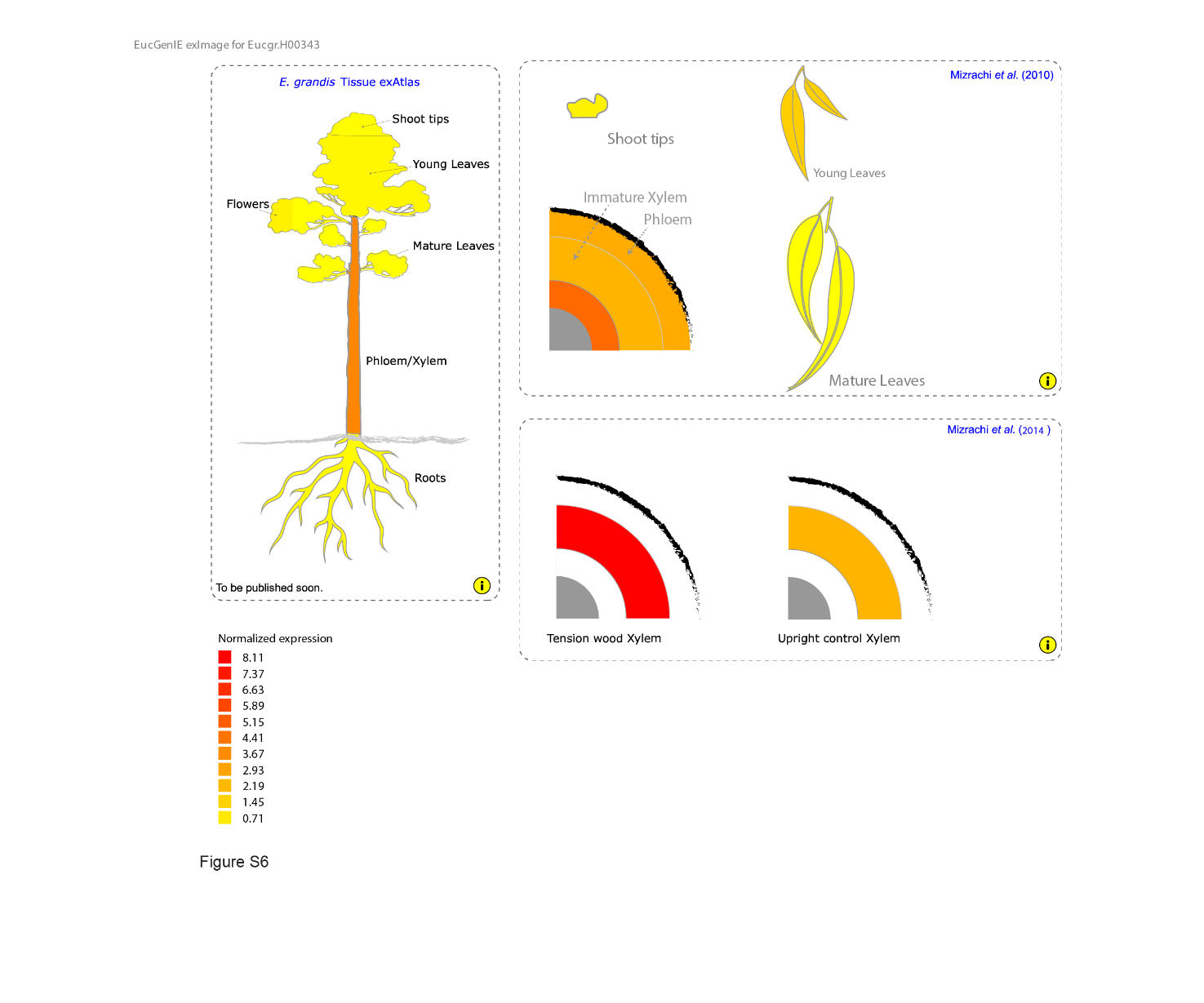

**Fig. S7. *Ed*XAPT and *Eg*XAPT coding sequences show high similarity.** The *Ed*XAPT coding sequence was amplified from *Eucalyptus dalrympleana* genomic DNA by PCR and sequenced by Sanger sequencing. The sequence was aligned to that of *Eg*XAPT using MUSCLE and translated using the ExPASy Translate Tool (https://web.expasy.org/translate/). Differing amino acids are highlighted in bold.

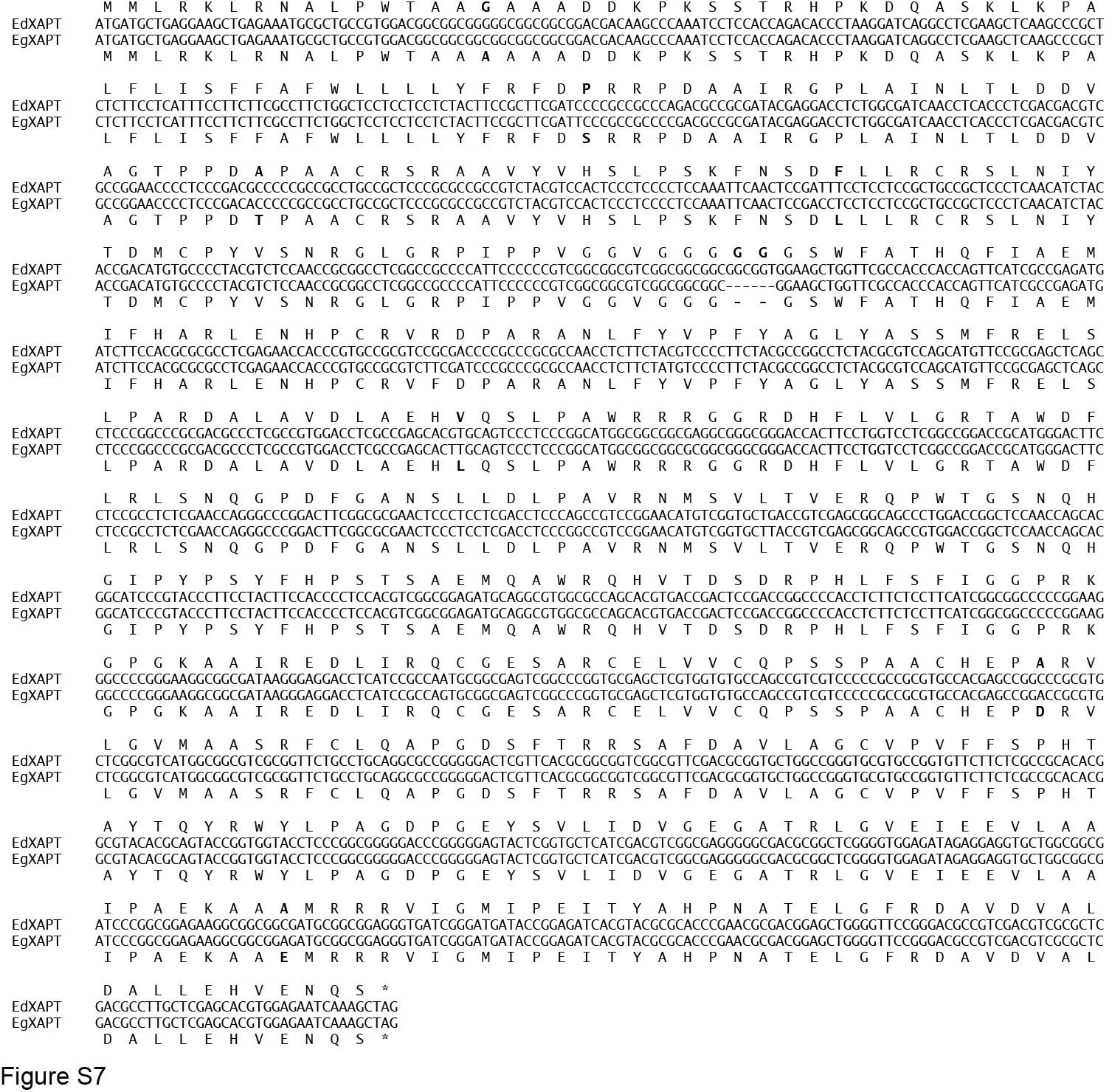

**Fig. S8. *Ed*XLPT and *Eg*XLPT coding sequences show high similarity.** The *Ed*XLPT coding sequence was amplified from *Eucalyptus dalrympleana* genomic DNA by PCR and sequenced by Sanger sequencing. The sequence was aligned to that of *Eg*XLPT using MUSCLE and translated using the ExPASy Translate Tool (https://web.expasy.org/translate/). Differing amino acids are highlighted in bold.

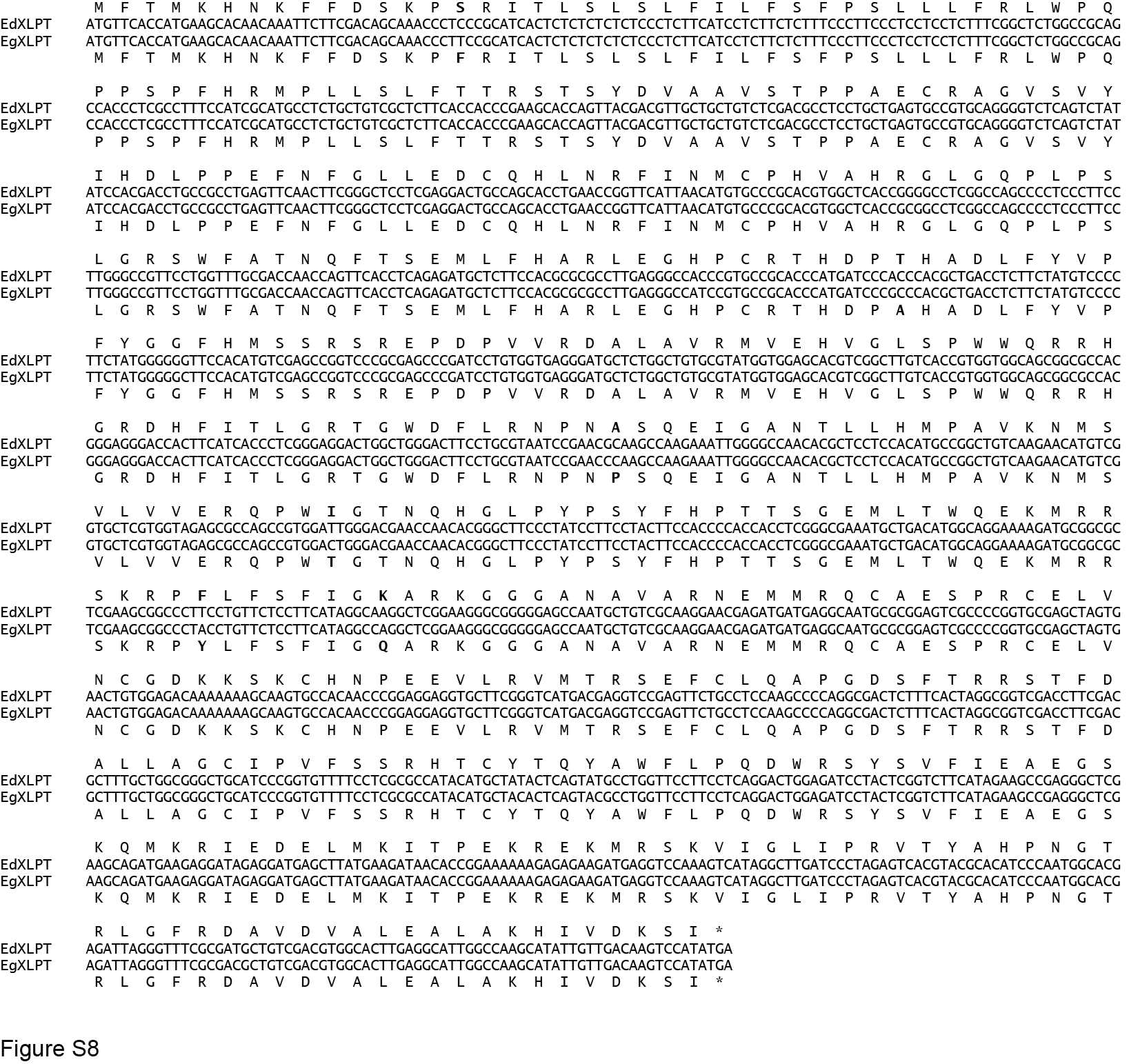

**Fig. S9. Other Myrtaceae plants contain U^Gal,[m]^ and U^Ara^*^p^*^,[m]^ structures in their xylan.** Xylem/pith tissue was extracted from five Myrtaceae plants by dissection before preparation of AIR. Alkali-extracted xylan was then digested simultaneously with *Np*Xyn11A xylanase and GH115 glucuronidase, which were subsequently removed by centrifugal filtration and ethanol precipitation. The resultant oligosaccharides were treated with or without AnGH35 β-galactosidase. β-GlcA'ase: β-glucuronidase. β-Gal'ase: β-galactosidase.

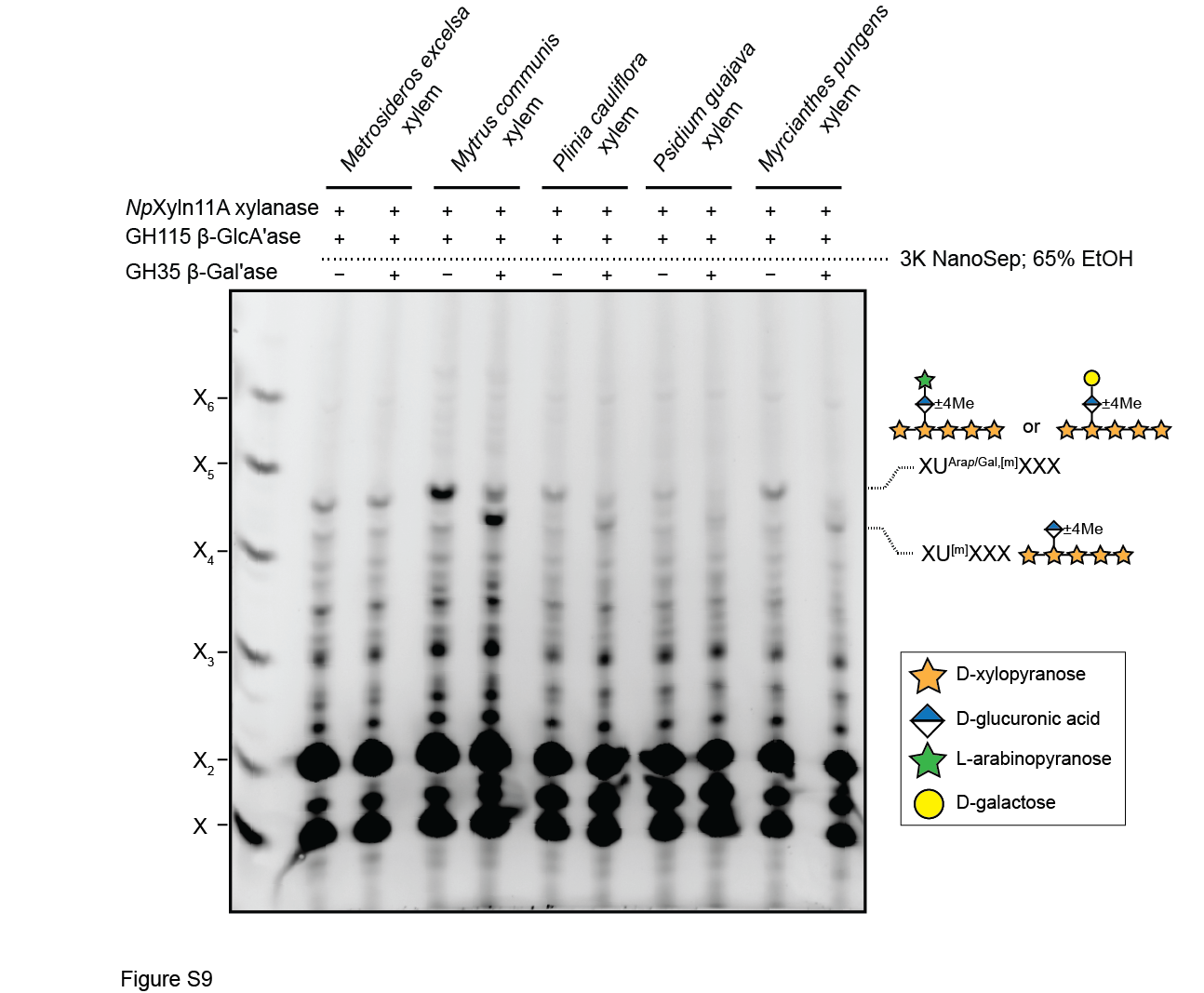

**Fig. S10. Other Myrtaceae plants contain U^Gal,[m]^ and U^Ara^*^p^*^,[m]^ structures in their xylan.** Cortex/phloem tissue was extracted from five Myrtaceae plants by dissection before preparation of AIR. Alkali-extracted xylan was then digested simultaneously with *Np*Xyn11A xylanase and GH115 glucuronidase, which were subsequently removed by centrifugal filtration and ethanol precipitation. The resultant oligosaccharides were treated with or without *An*GH35 β-galactosidase. β-GlcA'ase: β-glucuronidase. β-Gal'ase: β-galactosidase.

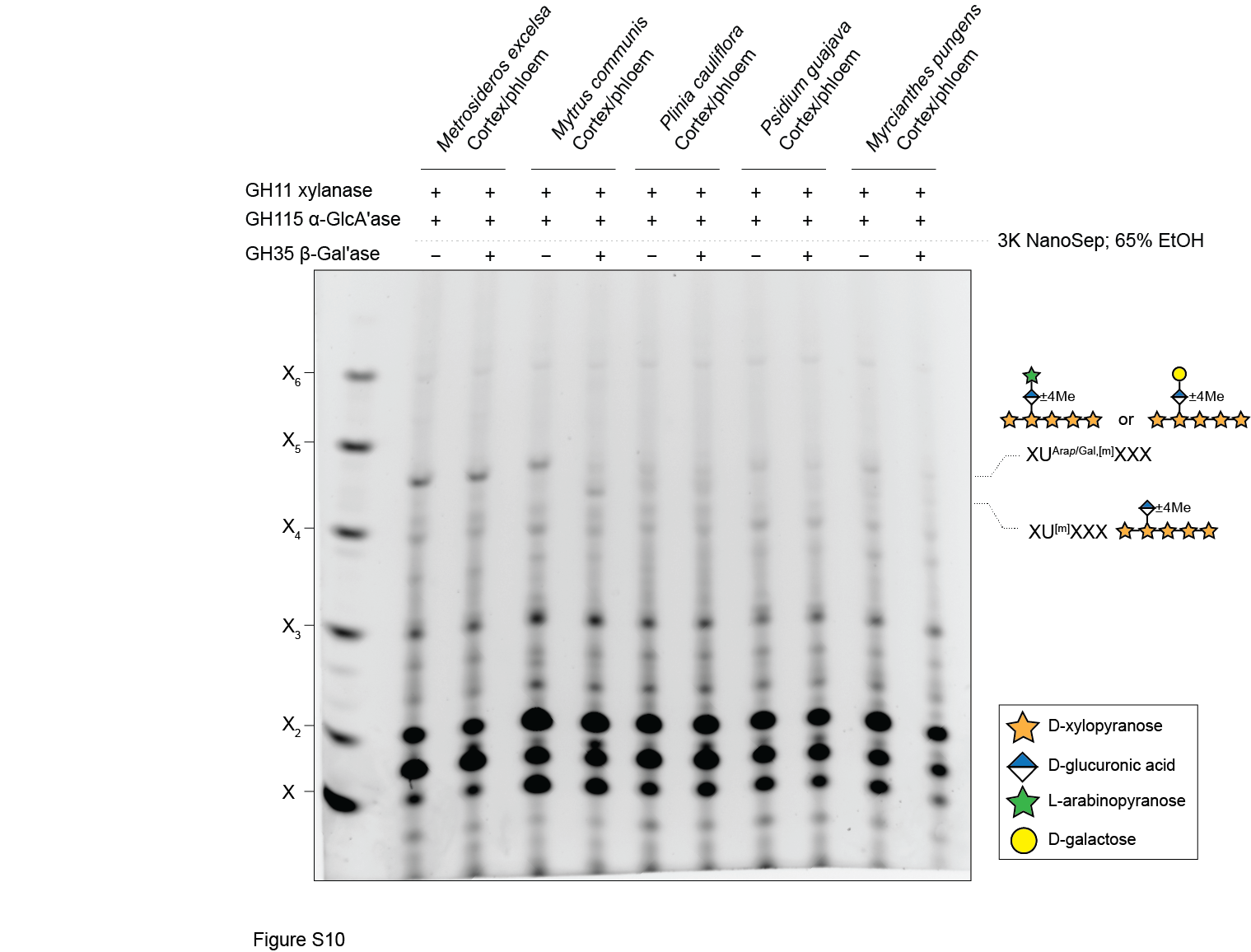

### Fig. S11. *xapt1* mutant plants show no obvious growth phenotype. Images show eight-week-old plants. Scale bar = 10 cm.

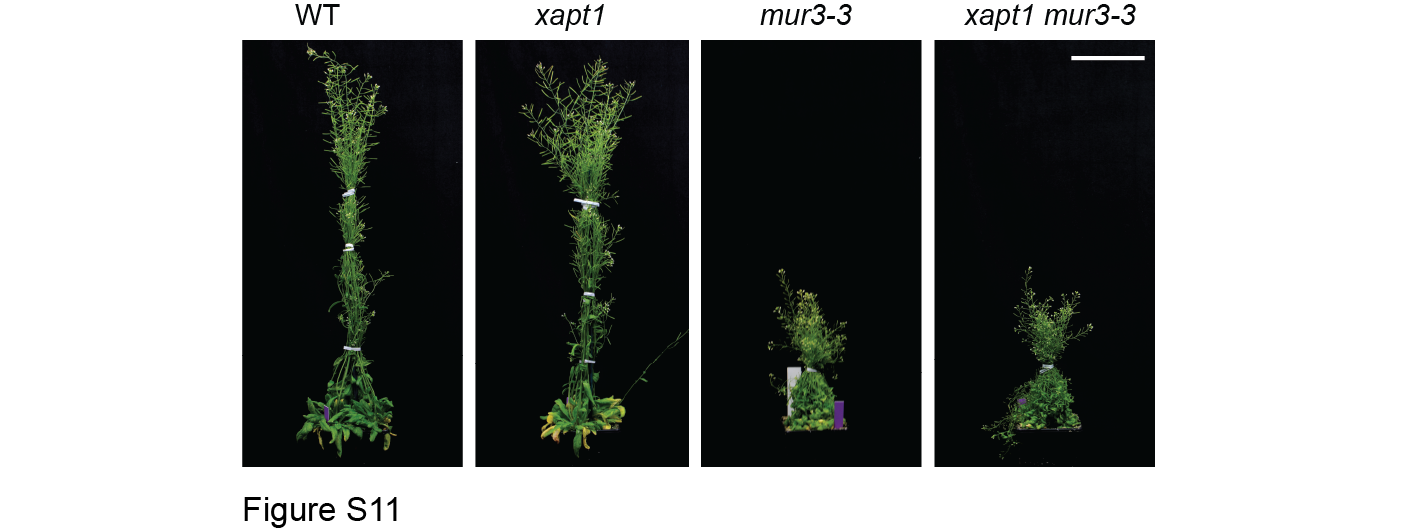

**Fig. S12. Xylan from WT bottom stem can be digested in the same way with *Ec*_D_Xyn30A, *Ec*_D_Xyn30A_Y255L_, *Ec*_P_Xyn30A, and *Ec*_P_Xyn30A_L255Y_.**

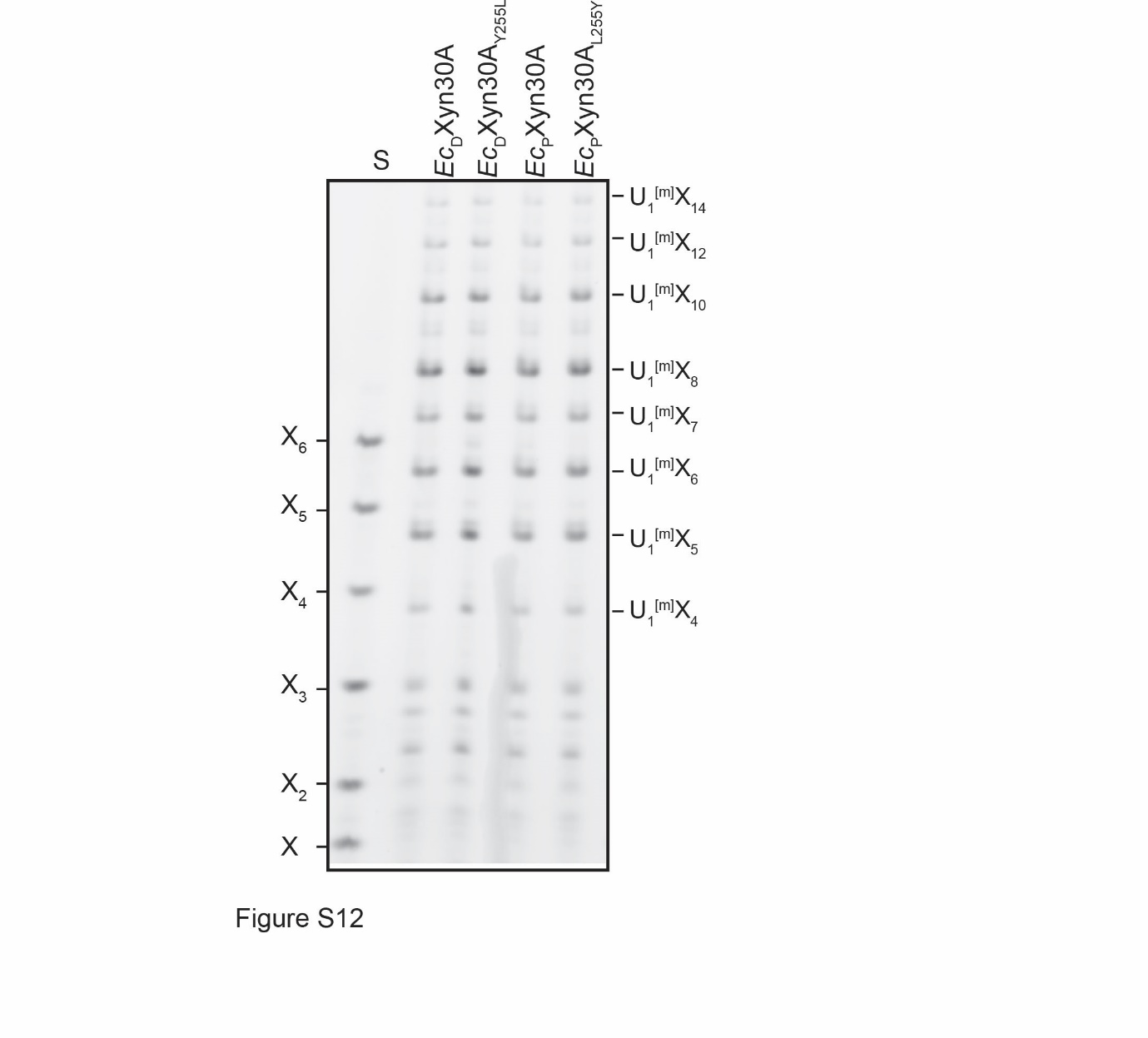

**Supplementary Tables**

**Table S1. Primers used in this study.**

| **Primer name** | **Sequence** |
| --- | --- |
| Primers for identification of T-DNA insertion mutant | |
| WiscDsLox437H04-LP | ACCACAATTTCACCGTCTACG |
| WiscDsLox437H04-RP | ATCGCATCGAATGTTGATCTC |
| salk_085915-LP | AGCGGGTGATATCATCATCTG |
| salk_085915-RP | TTGTTTATCTTTCACCACCGG |
| Salk_075020-LP | GGATTGAACAACCAATCATGG |
| Salk_075020-RP | TCACGGCGAACTTCTCTTATG |
| salk_057536-LP | TTGGAAAATGCAATTAAAGCG |
| salk_057536-RP | AGAAAACCGGTATACAACCCG |
| salk_053593-LP | TTTACAGGACGAGTTGGCG |
| salk_053593-RP | TCCCGTGACAAAGAAATGATC |
| salk_108349-LP | TCTCTGAGGACATTTCCGTTG |
| salk_108349-RP | TATGGTGGCGTACCTATTTCC |
| PCR primers used to amplify *Ed*XAPT and *Ed*XLPT | |
| *Ed*XAPT-LP | CACTATCTCCATTCCAGAAACCC |
| *Ed*XAPT-RP | CATCACATTACACCAACACAATC |
| *Ed*XLPT-LP | CTCACTTCTCCTCCATAACCC |
| *Ed*XLPT-RP | CAAGCTAATTGGATCGATCAAGC |
| PCR primers used to amplify GH30 enzymes | |
| *Ec_D_*Xyn30A forward | CTGTTGCTTTTAGTTCATCGATAGCATCAGCAGATACAGTCAAAATTGATGCG |
| *Ec_D_*Xyn30A reverse | GGTGATGGTGATGATGTTTGCCAACAAATGTTGTAAC |
| *Ec_D_*Xyn30A^Y255L^ forward | CAAACAAGTTTGGATGACAGAACATTATGTTGATTCAAAACAGAGCGCGAA |
| *Ec_D_*Xyn30A^Y255L^ reverse | ATGTTCTGTCATCCAAACTTGTTTGCCTGCATTTTGT |
| *Ec_P_*Xyn30A forward | CTGTTGCTTTTAGTTCATCGATAGCATCAGCAGACACAGTTAAGATCGACGC |
| *Ec_P_*Xyn30A reverse | GGTGATGGTGATGATGCTTAGAAACGAATGTAGTTACGC |
| *Ec_P_*Xyn30A^L255P^ forward | GCAGCTTTGGATGACTGAGCACCTTGTTGACTCTAAGCAATCAGCTAAC |
| *Ec_P_*Xyn30A^L255P^ reverse | GTGCTCAGTCATCCAAAGCTGCTTGCCAGCGTT |
